# Trajectories of hippocampal subregion development in the first years of life and their association with school-aged episodic memory outcomes

**DOI:** 10.64898/2026.05.12.724670

**Authors:** Sally M Stoyell, Jacob T Lundquist, Lana Hantzsch, Alexis Bunnell, Andrew Bunnell, Kathleen M Thomas, Damien A Fair, Brenden Tervo-Clemmens, Eric Feczko, Jed T Elison

## Abstract

Brain networks that support episodic memory development in the first years of life remain poorly understood. Protracted growth of regions such as the hippocampus have been suggested as a causal role in episodic memory development, but development of these memory brain networks and their role in episodic memory development is not yet fully elucidated. In this study, subcortical memory network regions (hippocampus, thalamus, amygdala) were segmented from MRI images in 835 visits spanning 0-4 years of age across 322 participants in the Baby Connectome Project. Hippocampal segmentations were further subdivided into head, body, and tail subregions manually for 426 visits, which were used to train models that automatically segmented hippocampal subregions for the remaining visits. 58 participants returned for an early school-age follow-up, including two episodic memory tasks. Volumetric growth trajectories differed across regions and across subregions within the hippocampus, with the head of the hippocampus showing steep growth that plateaued months later than the body or tail of the hippocampus. In the right hemisphere’s hippocampal head, age- and sex- adjusted volumes positively predicted future early school-age episodic memory performance. After accounting for total brain volume, the right thalamus also predicted memory performance. Total sleep duration at the follow-up visit accounted for performance variance above and beyond brain volume correlations. Altogether, results suggest that trajectories of growth and relationships between volume and episodic memory performance are region and subregion specific, and provide evidence for the important role of sleep in associations between brain networks and early episodic memory development.

**Significance:** The hippocampus is a critical structure in episodic memory, yet precise longitudinal developmental trajectories of this structure have yet to be elucidated. This study provides detailed, subregion specific hippocampal trajectories, and demonstrates that variation in these trajectories is associated with variation in later episodic memory performance. This insight fills a current gap in the literature delineating how brain development and episodic memory behaviors are related in the first five years of life. Considering this is the same age range during which adults begin to have long-term memories available from childhood, this gap represents an important opportunity to understand how changes in the brain support the development of basic episodic memory skills.

## Introduction

As a critical node in memory networks, understanding hippocampal development is critical in understanding the development of memory networks and their relation to episodic memory development. Studies show the hippocampus has a protracted growth period, with non-linear growth that is fastest in the first few years of life rather than just the first year (Gilmore et al., 2012; Holland et al., 2014; Utsunomiya et al., 1999). Within the hippocampus, it has been shown that there are different developmental trajectories between subregions (head, body, and tail) along a known long-axis gradient (Langnes et al., 2020; Nichols, Blumenthal, et al., 2023; Poppenk et al., 2013) across middle childhood (Canada et al., 2020). At a more granular level, there is evidence from primate literature that the subfields of the hippocampus develop at different times. CA1/CA2 fields, subiculum, and pre-subiculum develop earlier, while the dentate gyrus, and downstream fields such as CA3 develop later (Jabès & Nelson, 2015; Lavenex & Banta Lavenex, 2013). This differential timing of anatomical development suggests that the trisynaptic circuit in the hippocampus (Entorhinal cortex – dentate gyrus – CA3 – CA1) may demonstrate adult-like function later in development compared to the monosynaptic circuit (Entorhinal cortex – CA1) due to the protracted development in dentate gyrus and CA3 hippocampal subfields and connections to cortical regions (Bevandić et al., 2024).

Recent evidence suggests that even in infancy and toddlerhood, the hippocampus plays a direct role in memory function. Studies have shown hippocampal activation in response to statistical learning in infants as young as 3 months (Ellis et al., 2021), novel vs. familiar songs in 2 year olds (Mooney et al., 2021; Prabhakar et al., 2018), newly learned words in toddlers (Johnson et al., 2021), and most recently the encoding of photographs in infants as young as 12 months (Yates et al., 2025). As children enter early childhood, more aspects of episodic memory can be measured behaviorally. Studies relating the development of memory-related brain networks to episodic memory performance in childhood primarily focus on children older than 4 years. A meta-analysis of hippocampal development and memory ability broadly across childhood and adolescence showed a small positive association between hippocampal volume and memory performance (Botdorf et al., 2022). However, when studies looked specifically at narrower age ranges, age-specific hippocampal-memory associations were seen (Canada et al., 2019, 2021; DeMaster et al., 2014; Riggins et al., 2015, 2018). For example, in 4-6 year old children, larger volumes were correlated with better performance in the younger age group, while smaller volumes correlated with better performance in the older group (Riggins et al., 2018). While more studies are likely needed to confirm specific age-specific associations, these might reflect true developmental mechanisms such as synaptic pruning (Ghetti & Bunge, 2012). Associations also differed by subregion and subfield, suggesting differential developmental timing and specialization across the hippocampus (Bouyeure et al., 2021; J. K. Lee et al., 2020; Riggins et al., 2018; Schlichting et al., 2017; Tamnes et al., 2014; Vinci-Booher et al., 2023).

Early in development, much of brain development occurs during sleep, as infants and toddlers spend a good portion of their day sleeping. Ripple activity (high frequency oscillatory activity measured on EEG) in the hippocampus coordinates with sleep spindles (i.e., bursts of oscillatory activity between the thalamus and cortex) to replay memories and engage in memory consolidation during sleep (van den Berg et al., 2019). Numerous review articles have been written about the crucial role of sleep in the development of memory skills (Gómez & Edgin, 2016; Huber & Born, 2014). Both children and infants show better retention of declarative memories after a period of sleep as compared to a similar duration of wakefulness (Lokhandwala & Spencer, 2021; Seehagen et al., 2015), especially during slow wave sleep where sleep spindles are present (Kurdziel et al., 2013). Additionally, sleep duration has been shown to be correlated with both memory and hippocampal volumes in early childhood (Riggins & Spencer, 2020), highlighting the importance of studying the interplay between sleep, brain, and memory development.

To date, no studies in humans have examined detailed early life hippocampal trajectories at a subfield or subregion level. Even across the full hippocampus, sampling across ages is often too sparse to capture potential inflection points of growth occurring during the early years (Decker et al., 2020; Dick et al., 2022; Gilmore et al., 2012; Li et al., 2023; Uematsu et al., 2012; Utsunomiya et al., 1999). While it is known that the hippocampus undergoes a longer period of growth relative to other brain regions, the shape of growth trajectories across the hippocampus, including specific non-linear trends, is still unknown. To date, studies of memory network development in early childhood have used primarily cross-sectional data and have focused primarily on children 4 years of age or older, representing an important gap in our understanding of early relations between brain development and memory behaviors. Here we define subregion-specific trajectories in what is currently the most densely sampled study across the first years of life, providing important nuance to previous examinations of subcortical and hippocampal growth patterns. We further characterize subregion-specific trajectories by their association with early school-age outcomes, including sleep and episodic memory performance.

## Materials and Methods

### Participants

Data were used from the Baby Connectome Project (BCP), a study of infants and young children 0-60 months old that measured brain and behavioral development across the first 5 years of life. Details on study design, recruitment, and overall methodology have been described previously (Howell et al., 2019). Briefly, infants were recruited from research participant registries based on both state-wide birth records and outreach to the broader communities around the University of North Carolina at Chapel Hill (UNC) and the University of Minnesota (UMN). Infants were eligible for the BCP if they 1) were born between 37–42 weeks gestation, 2) had a birth weight appropriate for gestational age, and 3) had no major pregnancy or delivery complications. Participants were excluded if they: 1) were adopted, 2) had a first degree relative with autism, intellectual disability, schizophrenia, or bipolar disorder, 3) had any significant medical and/or genetic conditions affecting growth, development, or cognition, 4) had any contraindication to MRI, or 5) had caregivers who were unable to communicate in English at a level to provide informed consent. All procedures were approved by the UNC and UMN Institutional Review Boards. Data from this cohort were collected via an accelerated longitudinal design, with infants completing up to nine neuroimaging and behavioral sessions, and most infants completing between 1-6 visits.

See Figure 1 for a flowchart of BCP MRI visits. In total, 1233 MRI visits were attempted across 470 participants in the BCP study. 897 visits across 364 participants yielded quality structural MRI images and were successfully run through structural processing pipelines. Participant ages and other demographic characteristics can be seen in Figure 2 and Table 1. Ages ranged from 0-5 years old, with more dense sampling of visits prior to 2 years. Race and ethnicity of the sample matched the census data for the metro areas around UMN and UNC for each site respectively. 23 visits failed segmentation quality checks, including one participant with an incidental clinical finding on MRI. These visits were removed from further analyses. Based on the sparseness of the data after 45 months of age, analyses were limited to visits less than 45 months of age, for a total of 835 visits.

**Figure 1.**
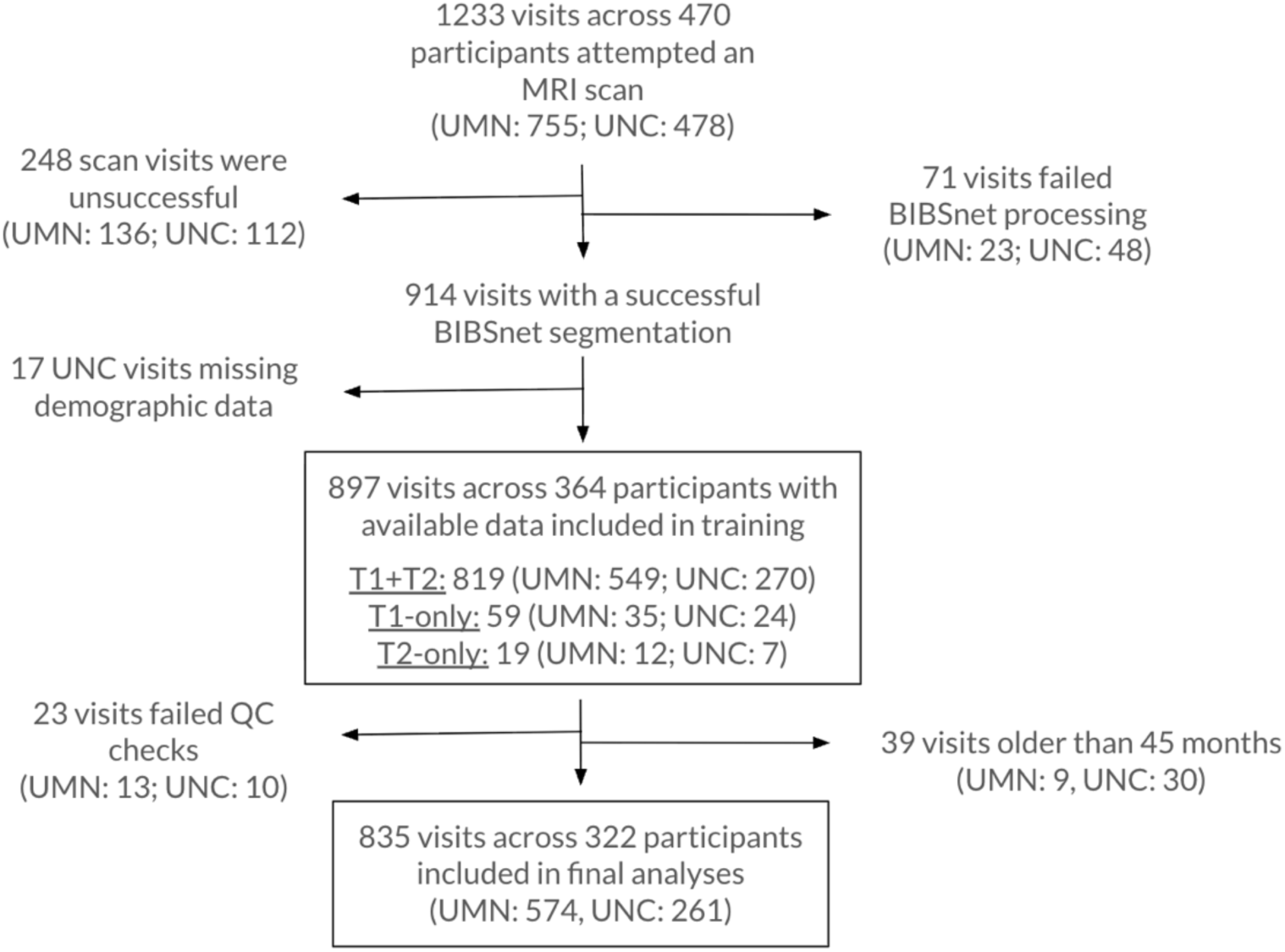
Flowchart of participants and visits from the BCP study.

**Figure 2.**
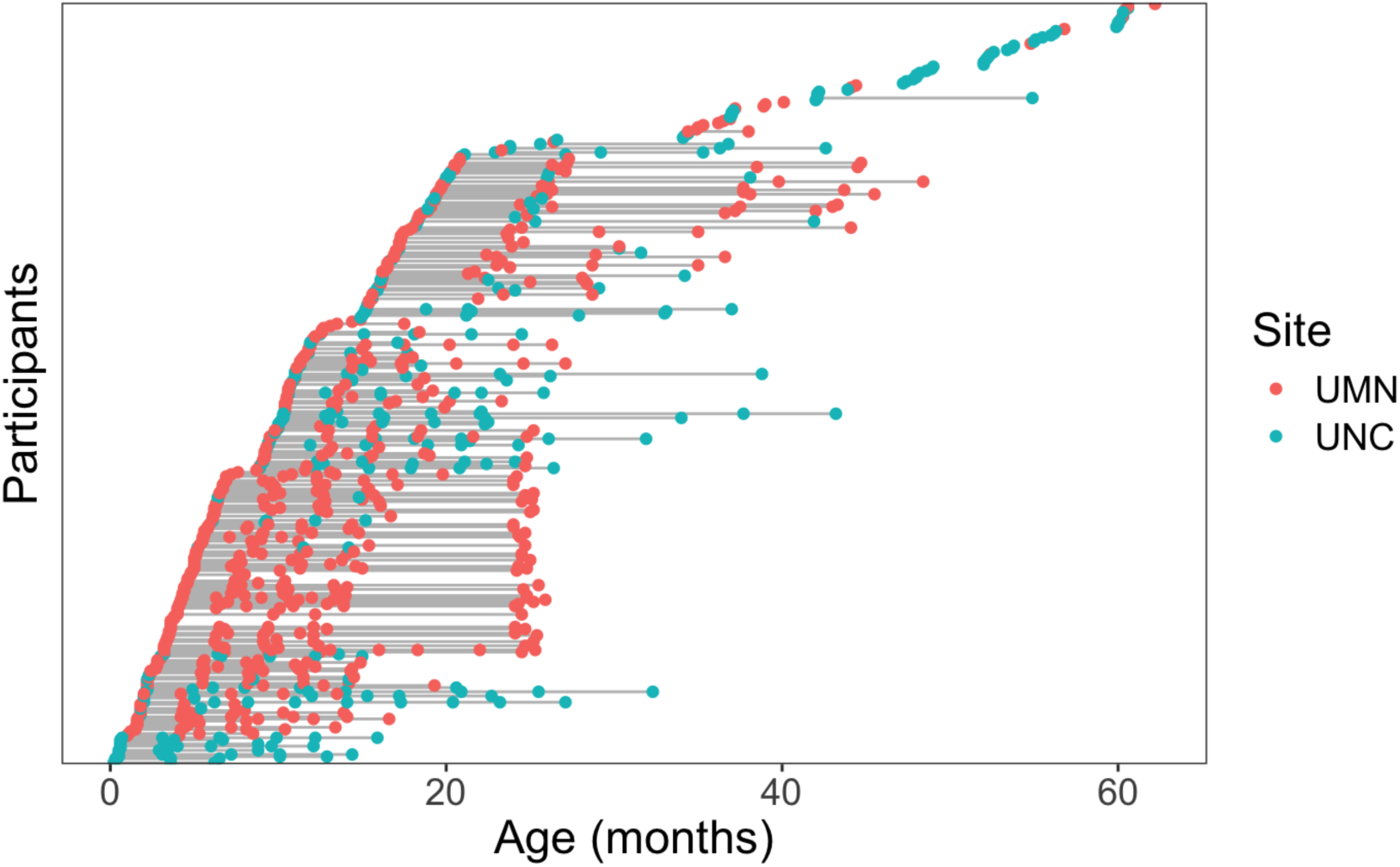
BCP participants with structural MRI data.

**Table 1.**
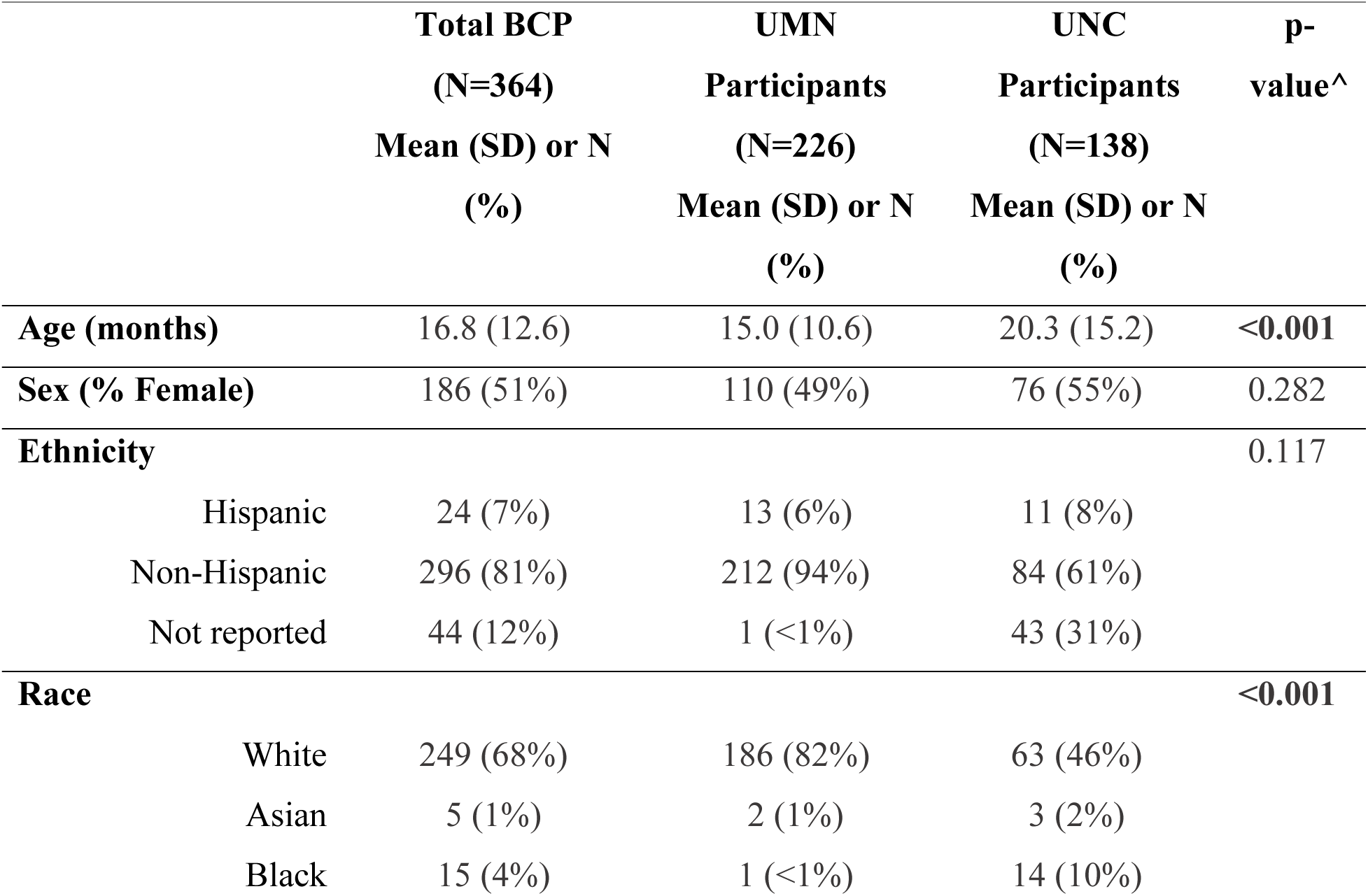
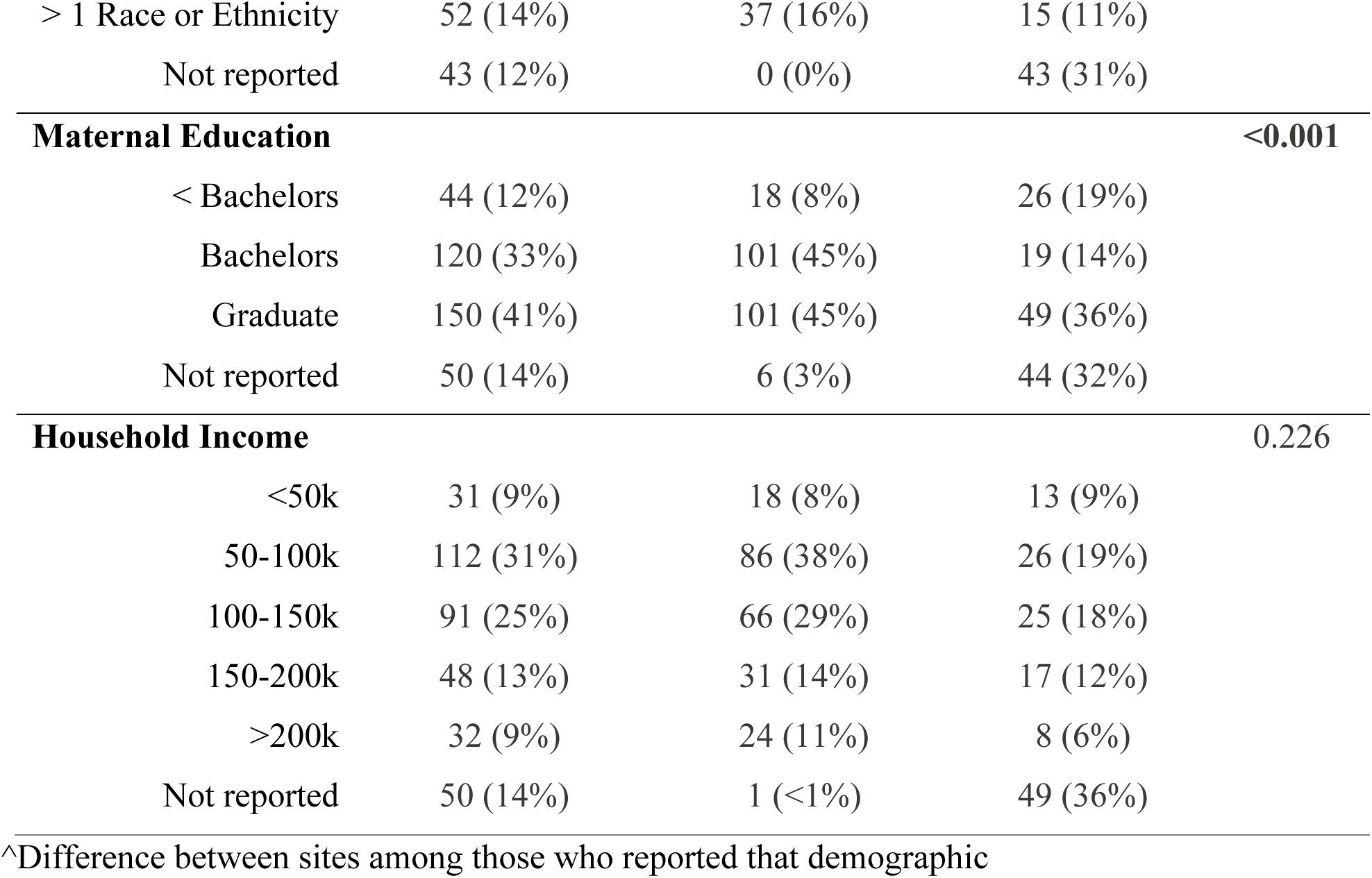
BCP Participant Demographics.

To measure memory specific outcomes, two separate episodic memory tasks were added to an early-school-age follow-up of BCP participants at the University of Minnesota site. For this subset, BCP participants with successful imaging or eye-tracking data at one of their visits between 0-5 years were sent invitations to complete a school-aged survey. Those that completed the survey (N=121) were invited for two visits to the university for in-person behavioral testing including episodic memory follow-up (N=58). Participants ranged between 3-7 years of age, but 91% were between 4-6 years (Mean=5.5, SD=0.93). All but one of those participants had at least one successful structural MRI scan from an earlier BCP imaging visit between 0-5 years. No differences were found in demographic characteristics (Table 2) for this subsample relative to UMN BCP imaging participants who did not return for a school-aged memory visit.

**Table 2.**
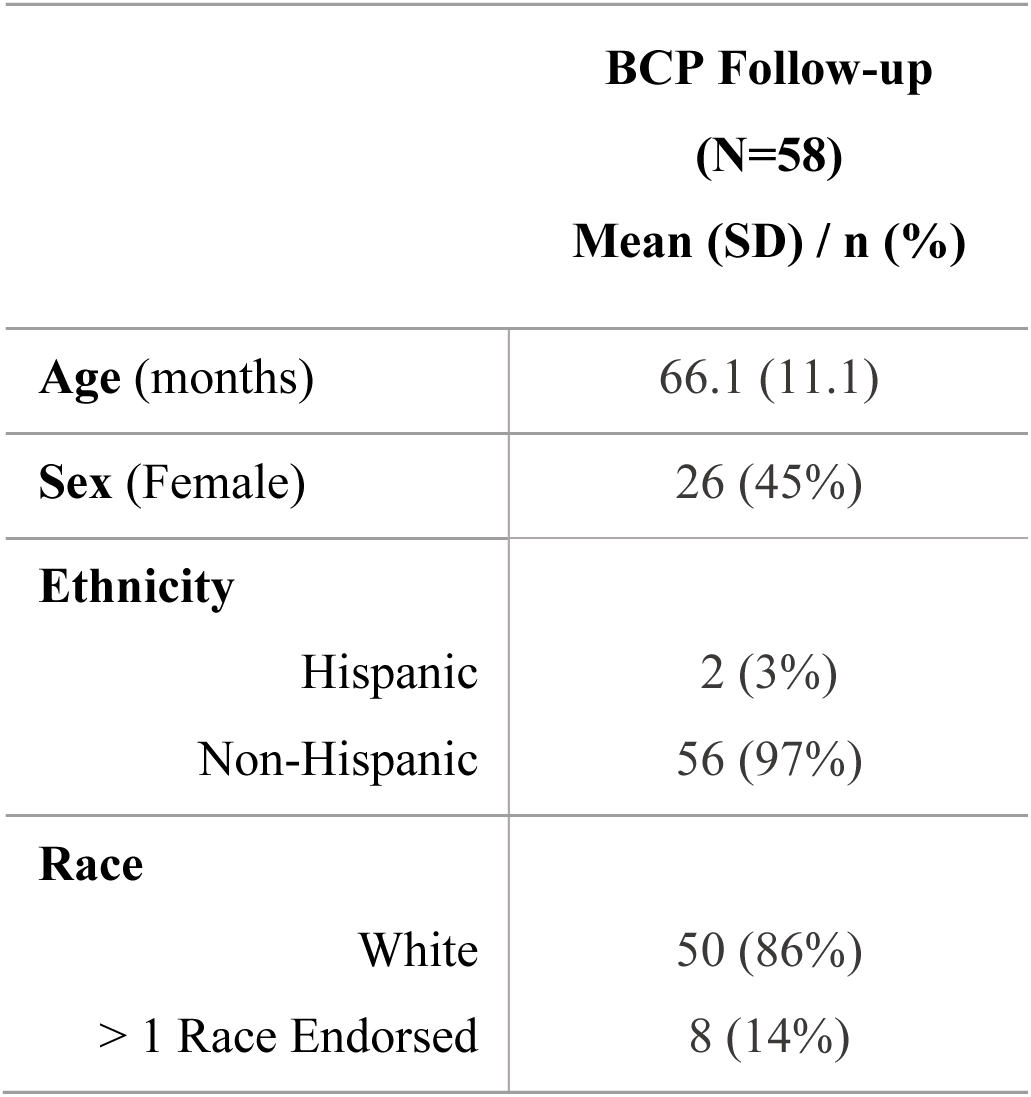
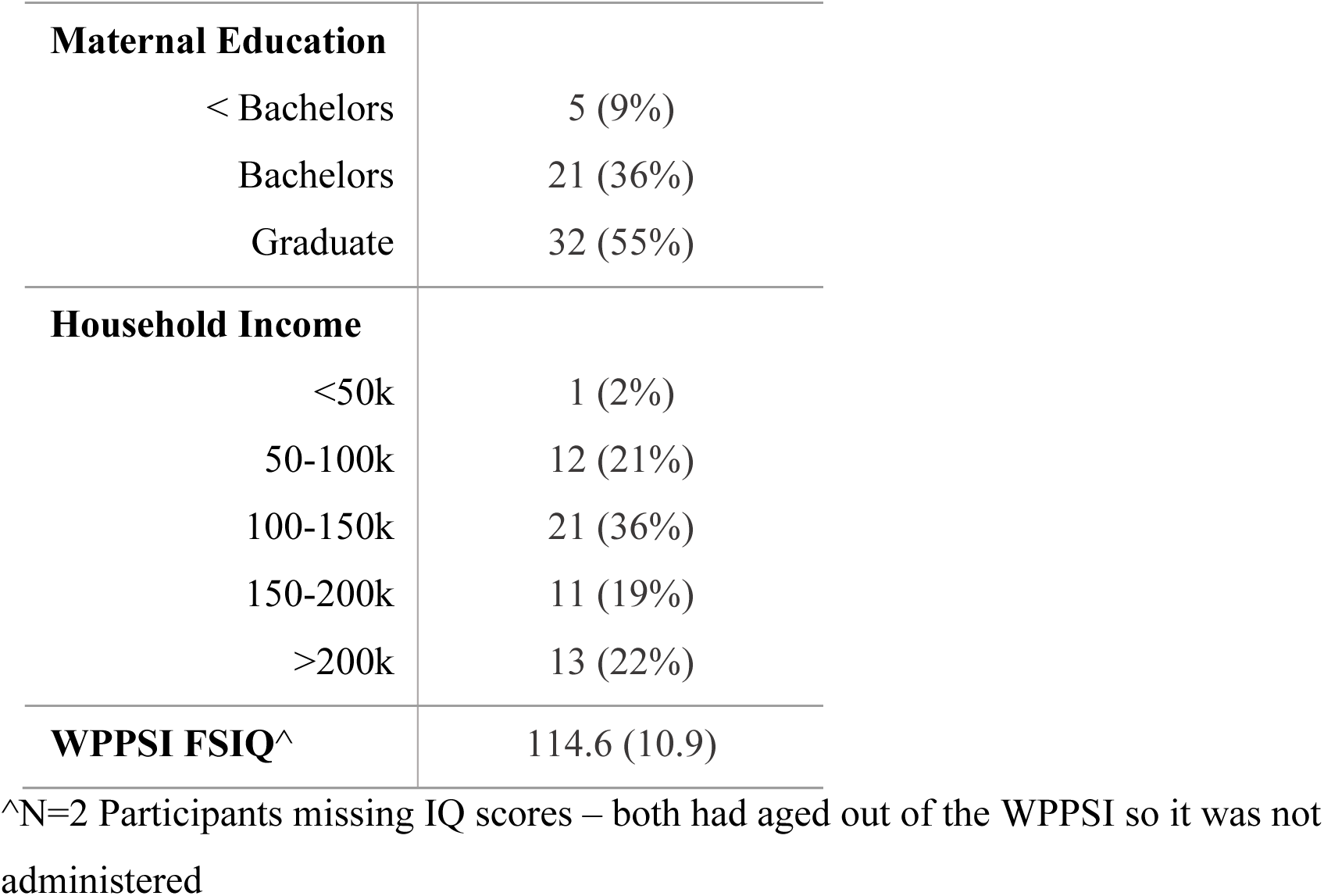
BCP School-age Follow-up Participant Demographics.

| **BCP Follow-up**<br>**(N=58)**<br>**Mean (SD) / n (%)** |  |
| --- | --- |
| **Age (months)** | 66.1 (11.1) |
| **Sex (Female)** | 26 (45%) |
| **Ethnicity** |  |
| Hispanic | 2 (3%) |
| Non-Hispanic | 56 (97%) |
| **Race** |  |
| White | 50 (86%) |
| > 1 Race Endorsed | 8 (14%) |
| **Maternal Education** |  |
| < Bachelors | 5 (9%) |
| Bachelors | 21 (36%) |
| Graduate | 32 (55%) |
| **Household Income** |  |
| <50k | 1 (2%) |
| 50-100k | 12 (21%) |
| 100-150k | 21 (36%) |
| 150-200k | 11 (19%) |
| >200k | 13 (22%) |
| **WPPSI FSIQ<sup>^</sup>** | 114.6 (10.9) |

<sup>^</sup>N=2 Participants missing IQ scores – both had aged out of the WPPSI so it was not administered

### Imaging

State-of-the-art MRI imaging acquisition techniques were used with modifications to account for the unique challenges of scanning infants. Images were collected during natural sleep on a 3T Siemens Prisma scanner using a 32-channel head coil. Structural T1w and T2w images were collected from all infants. T1w images were collected using an MPRAGE sequence: TR 2400ms, TE 2.24ms, TI 1600ms, Flip angle 8°, and resolution = 0.8 × 0.8 × 0.8 mm3. T2w images were collected using turbo spin-echo sequences: Turbo factor 314, Echo train length 1166ms, TR 3200ms, TE 564ms, resolution = 0.8 × 0.8 × 0.8 mm3, with a variable flip angle. These sequences were collected as part of the broader BCP protocol (Howell et al., 2019) which included additional images, such as diffusion weighted images, that are beyond the scope of this paper.

### Behavioral Tasks

#### Source Memory Task

A source memory task that has previously been used to measure episodic memory in preschool children was adapted for use in this study (Callow et al., 2020; Riggins et al., 2018). Children were taught 12 age-appropriate facts (ex. “A group of rhinos is called a crash!”) from two different sources: 6 from a video of a girl “Abby” and 6 from a video of a puppet “Henry”. Before each fact was taught, children were asked if they already knew the answer to the fact (ex. “What is a group of rhinos called?”). If children already knew the fact, an additional novel fact was taught, such that all children learned 12 *novel* facts. Participants were counterbalanced across three study parameters: which of three different lists of facts was presented, the order in which the two source videos were played (Abby or Henry first), and the portion of the fact list that Abby or Henry taught. Children were instructed to learn the facts and were told they would earn a prize if they remembered enough facts later. Children were not told that they would be expected to recall the source of the fact, such that the encoding of the source of the fact was incidental. After a one-week delay, children were asked a set of trivia questions. Children were instructed that some of the questions would be from the videos they watched the week before, and some would be things that they might have learned from somewhere else. Children were told they should ask for a hint if they did not know an answer. Questions were either facts learned in the previously presented videos, facts that children would be expected to know (ex. “What color is grass?”), or facts that young children would likely not know and had not been taught in the session (ex. “What is the colored part of your eye called?”). If children indicated that they did not know or asked for a hint, multiple choice options were provided, and children were asked to select one of the offered options. After responding to the trivia question, children were asked where they had learned that fact. Children were prompted at the recall with still images of Abby and Henry, as well as icons representing other potential source options (parent, teacher, guessed, just knew it). If a child indicated they didn’t know where they learned the fact, they were asked if someone taught it to them or if they just knew it/guessed. If they indicated that someone taught them, but that they didn’t know who, they were explicitly presented with the multiple-choice options (parent, teacher, person on the video, puppet on the video, just knew/guessed). The still image prompts were implemented part way into the study, such that the first 16 participants did not receive these prompts. As such, sensitivity analyses were completed to determine whether the additional cues impacted results.

Of the 58 participants, 3 could not return for a second visit, so the source memory task was not attempted. Of the 55 participants who attempted the source memory task, 1 was excluded due to experimenter error during recall. Participants averaged 7.4 days between encoding and recall, with 83% of the sample returning between 6-8 days later, and only one participant beyond 14 days. No correlation was seen in the outcome of interest and the number of days between encoding and recall. Four participants had experimenter error on one or two specific questions at recall and so outcomes were calculated out of 10 or 11 novel facts learned instead of 12. Due to this, outcomes of interest were calculated as proportions of facts remembered rather than total number of remembered facts.

The main outcome of interest for this source memory task was the proportion of taught facts where the participant recalled or recognized both the correct answer and attributed it to the correct source (Abby or Henry). Exploratory analyses examined the type of source errors made, including the proportion of sources incorrectly attributed to sources outside the experiment (“Other”), rather than attributing it to guessing/just knowing (“Guess/Know”) or correctly attributing it to the experiment but the other source (“Intra-Experimental”).

### Mnemonic Similarities Task (MST)

A pattern discrimination task designed to measure episodic memory in preschool children was also included in this study (Bouyeure et al., 2021; Canada et al., 2019). This task consisted of an incidental encoding phase, a training phase, and a recall phase. In the incidental encoding phase, children made “inside” vs. “outside” judgments for pictures of objects by pressing one of two buttons that said “inside” or “outside” audibly when pressed. “Inside” and “outside” judgements were not of interest and were used only to keep children engaged. Images were displayed on the screen for a fixed duration of 2 seconds. In the training phase, children were taught to categorize images as “exactly the same” as each other, “kind of the same” as each other, or completely different from one another. Once children demonstrated that they understood these concepts, they were shown a new set of images and asked to label each as “exactly the same” (Same; Targets), “kind of the same” (Similar; Lures), or “new picture” (New; Foils) compared to images viewed in the “inside/outside game” (Figure 3). Children provided responses on three talking buttons, which were then recorded by the experimenter using the keyboard. No time constraints were placed on the responses. Children were counterbalanced into one of 6 sets of images, and the order of images in both the encoding and recall phases was randomized for each participant. All images were presented in Eprime 3.0. All 58 participants attempted the MST task. Of those, one failed to complete the task, for a final sample of 57 participants with MST data.

**Figure 3.**
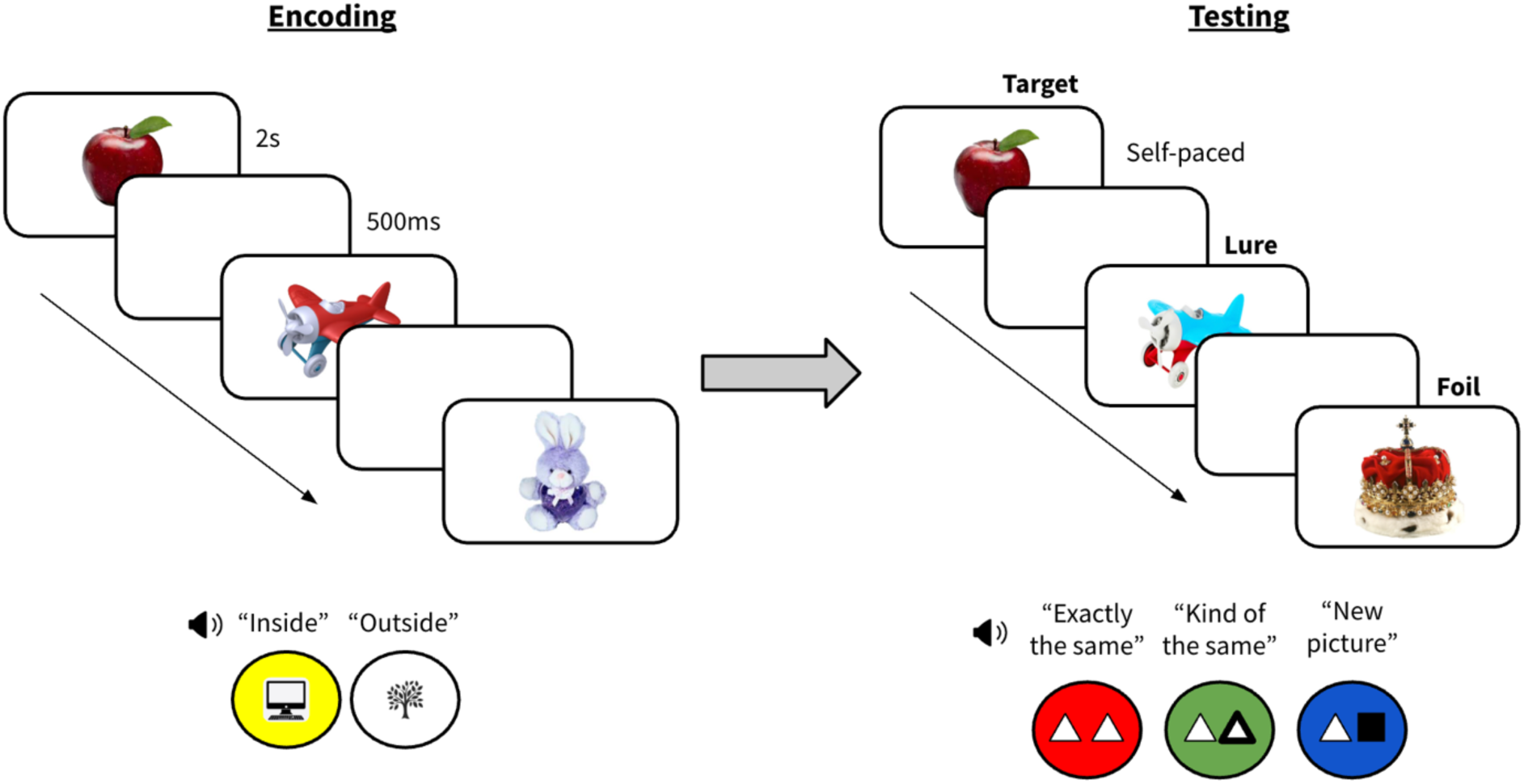
Figure depicting the setup of the MST.

For the MST, the main outcome of interest was a lure discrimination index calculated by subtracting the proportion of “exactly the same” responses to images that were similar but not identical (lures) from correctly assigned “kind of the same” responses given to similar items (i.e. similar|similar - same|similar) (Ngo et al., 2018). This index ranges from -1 to 1, where positive values indicate successful pattern discrimination, negative values indicate overgeneralization, and near zero values indicate performance close to chance.

A recognition score was calculated as the proportion of “same” trials correctly assigned “same” minus the proportion of “same” trials incorrectly assigned “new” (i.e. same|same - new|same), essentially measuring how well a child remembered the target images. This averaged 0.74 across all participants, ranging from 0 to 1 across the full range of possible values. As most children performed well for “same” and “new” images overall, for sensitivity analyses, participants were only included if their overall performance reached a 0.4 on this recognition score (Supplemental Figure 1). This “recognition filter” was used as a proxy of the child’s understanding and engagement with the task and is similar to general item accuracy measures used in the literature (Canada et al., 2019; Ngo et al., 2018; Stark et al., 2019). Eight participants were excluded when this filter was applied for sensitivity analyses.

#### Wechsler Preschool and Primary Scale of Intelligence (WPPSI)

Participants also completed the WPPSI, a standardized measure of general cognitive abilities. Block Design, Information, Matrix Reasoning, Bug Search, Picture Memory, and Similarities subtests were used to calculate a full-scale IQ composite (FSIQ).

### Parent Questionnaires

Among other questionnaires, parents completed two assessments of their child’s sleep habits. Throughout the original BCP data collection visits, parents completed the Brief Infant Sleep Questionnaire (BISQ). The BISQ is a validated parent questionnaire that assesses sleep behaviors in infants (Sadeh, 2004). The two measures of interest from the BISQ were the total sleep duration and the sleep problems question, which is a single question asking the parent if they believe their child’s sleep to be a problem. At the school age follow-up, parents completed the Children’s Sleep Habits Questionnaire (CSHQ). Similar to the BISQ, the CSHQ is a validated parent questionnaire that assesses sleep behaviors in children. (Owens et al., 2000). The two measures of interest from the CHSQ were the total sleep duration and the total sleep disturbances score, a summed score derived from 33 questions about sleep problems that a child could potentially experience.

### Image Processing

T1 and T2 images from each available BCP session were run through BIBSnet (Hendrickson et al., 2025), a pipeline that utilizes nnU-Net (a deep-learning based learning software) and high-quality training data (Feczko et al., 2025) to create anatomical MRI segmentations (https://github.com/DCAN-Labs/BIBSnet). Hippocampal segmentations were also generated by this pipeline. The hippocampus was segmented further into head, body, and tail regions, allowing us to analyze the developmental trajectories of these subregions separately. While segmenting into subfields along histological boundaries (e.g., CA1/CA2, CA3, dentate gyrus) would be even more accurate, the resolution of the BCP anatomical data did not support this level of segmentation, which usually requires high-resolution scans specifically of the medial temporal lobe. For 426 visits across 169 participants, subregion segmentation was completed manually by three coders. Based on the availability of the pipelines at the time of segmentation, coders either started with BIBSnet version 3.2 or 3.4. Coders followed a standard protocol used in studies of early childhood hippocampal development (Riggins et al., 2015). Boundaries were defined in coronal slices, with the boundary between the head and body defined as the uncal apex and the boundary between the body and tail defined as the slice where the fornix separates from the hippocampus. Coders worked in ITK-SNAP (https://www.itksnap.org; (Yushkevich et al., 2006)), using both the T1 and T2 images when both were available, to subdivide the existing hippocampal segmentation into head, body, and tail regions and edit the overall segmentation as necessary. BIBSnet model starting points were trained on data that did not include the tail in the definition of the hippocampus, so the coders often needed to manually identify portions of the tail.

Dice scores were calculated as inter-rater reliability metrics between the three coders for 10 visits from each of the BIBSnet models (3.2 or 3.4), for a total of 20 visits. Inter-rater reliability metrics for hippocampal head, body, and tail segmentation for both BIBSnet models (3.2 or 3.4) were very similar, indicating that coders were able to create similar outputs from both models. Dice scores for the left and right head and body were all above 0.85, with an average dice score across raters and across visits greater than 0.95 for each subregion for each model. Dice scores for the tail were slightly lower, averaging 0.78 for both models. As the output from both BIBSnet models excluded the tail of the hippocampus, coders had to manually draw the tail, while the head and body regions of the hippocampus required only manual edits of any segmentation errors and then identification of the boundary between the head and body. Thus, it is not surprising that dice scores for the manually drawn tail would be lower than for the head and body regions. The dice scores for the tail of the hippocampus fell just below a typical “good” cut-off of 0.8, suggesting that the head and body reliability can be considered “very good”, and the tail reliability can be considered “acceptable” to “good”. These dice scores are all at or above average interrater reliability dice metrics used in subcortical structure segmentation for other model training data (Wang et al., 2022).

For the remaining 464 visits across 234 participants, hippocampal subregions were segmented automatically. There were no differences by segmentation method (manual or automatic) in age (median_manual_ = 13.5 months, median_auto_ = 13.4 months, Wilcoxon signed-rank test, W=101,778, p-value = 0.442), sex (x²=0.783, p=0.376), or site (UMN vs. UNC, x²=3.181, p=0.075). Participant visits with both T1 and T2 images available were prioritized for manual segmentations, as were participants who completed more than one visit, which may explain the trend towards a significant difference by site.

To train the automatic segmentation model, visits with manually segmented hippocampal subregions were divided into training and test sets. Participants were excluded from the training-test sets if they had follow-up memory outcome data (N=58), if they had a known clinical finding from imaging scans (N=1), or if the visit did not have both a T1 and a T2 image. In total, 282 visits across 113 participants contributed to the train-test split, with 227 visits across 103 participants in the training set and 55 visits across 48 participants in the test set. Visits were stratified across the train-test split by age, sex, model, and site. The training set was used as input data for an nnUnet-based deep learning model. For the nnUnet-based deep learning model, the model was trained using a GUI-based version (https://github.com/DCAN-Labs/dcan-nn-unet) of nnUnet (Isensee et al., 2021), that incorporates SynthSeg (Billot et al., 2023), which creates synthetic images to bolster the generalizability of the training data. The model was trained on T1/T2 images aligned in the same space, as output by BIBSnet, to reduce the risk of model performance issues related to alignment. Final segmentations were transformed back into native space using the reverse transform provided by BIBSnet. A preexisting hippocampal segmentation software, ASHS (Yushkevich et al., 2015), was also trained but showed lower quality segmentations than the nnUnet model (See Supplemental Analyses).

Amygdala and thalamus volumes were also extracted directly from the BIBSNet pipeline, as additional subcortical regions known to contribute to broader memory networks.

### Experimental Design and Statistical Analysis

Generalized additive mixed models (GAMMs) were used to estimate trajectories of growth in each brain region of interest. GAMMs use penalized splines to quantitatively define potentially non-linear developmental trajectories. Models included a smoothed effect of age, a fixed effect of sex, a fixed effect of site, and a per-participant random intercept. More complex models were tested, including an additional random slope per participant, but did not converge across all brain regions and had qualitatively and quantitatively highly similar results. Thus, to balance model complexity and parsimony and to ensure consistency across brain regions, the model including only the random intercept was chosen. Models were run both with and without total brain volume (TBV) as a covariate to determine both absolute and relative growth rates as suggested by recent best practices in structural brain development analyses (Vijayakumar et al., 2018). All resulting developmental trajectories were inspected with consideration to biological and developmental plausibility. Initial models in the thalamus yielded overly complex smooths with excessive curvature that were unlikely to reflect biologically plausible patterns. Thus, initial basis dimensions (*k*) were decreased to *k*=6 to prevent overfitting until smooths demonstrated biologically plausible trajectories. Finite derivatives were calculated using the *gratia* package in R (Simpson, 2024) in 1/10^th^ month increments. Simultaneous 95% CI of the derivatives were calculated using posterior simulation (with 10,000 iterations), and growth was deemed statistically significant in a region over the ages in which the 95% CI of the derivative did not cross zero (p<0.05) (Tervo-Clemmens et al., 2023).

To compare brain development measures with episodic memory outcome, residuals from each of the GAMMs trajectories were calculated for all visits with participants who had follow-up memory measures. This is akin to normative modelling, such that we determined how different a participant’s regional brain volume was from their predicted volume based on their age and sex. Separate models were created for MST pattern discrimination and source memory task outcomes, which measure unique aspects of episodic memory. Each model included the brain regions of interest, age at MRI, age at memory task, and the number of visits that a participant contributed. Since there was only one memory outcome score per participant, but multiple brain imaging assessments, the number of visits each participant contributed was used to weight participants by their number of visits.

Additional models comparing brain development measures with episodic memory outcomes used the GAMMs residuals that accounted for total brain volume to understand how relative brain region size related to behavioral outcomes. An additional model included FSIQ to understand the effects of regional brain volumes on memory outcomes above and beyond the effects of FSIQ.

Finally, to determine the effect of sleep behaviors on associations between brain development and memory performance, models including sleep measures from the BISQ and CHSQ were also tested. The two CHSQ sleep measures were each added as an additional covariate along with the GAMMs residual to determine if concurrent sleep behaviors correlate with episodic memory performance above and beyond any effect of early regional brain volumes. BISQ measures were added to the model predicting episodic memory performance to test if sleep behaviors during development predict additional variation in later episodic memory performance above and beyond any effect of the early brain measures.

## Results

### nnUnet Hippocampal Subregion Segmentations

Dice scores comparing the overlap between the nnUnet automatic segmentations and the test set of the manual segmentations indicated a high degree of agreement. The mean dice scores across all head and body regions were >0.89. The left and right tail respectively had a mean dice score of 0.8 and 0.78, hovering just on the line of “good”. This is likely due to the wider variability in the underlying manual segmentations used in the training data set, as the tails had to be fully drawn manually. Hippocampal subregion segmentation trajectories looked essentially identical between manual and nnUnet automatic segmentations (Figure 4), which supported the decision to collapse all data across manual and nnUnet automatic segmentation approaches.

**Figure 4.**
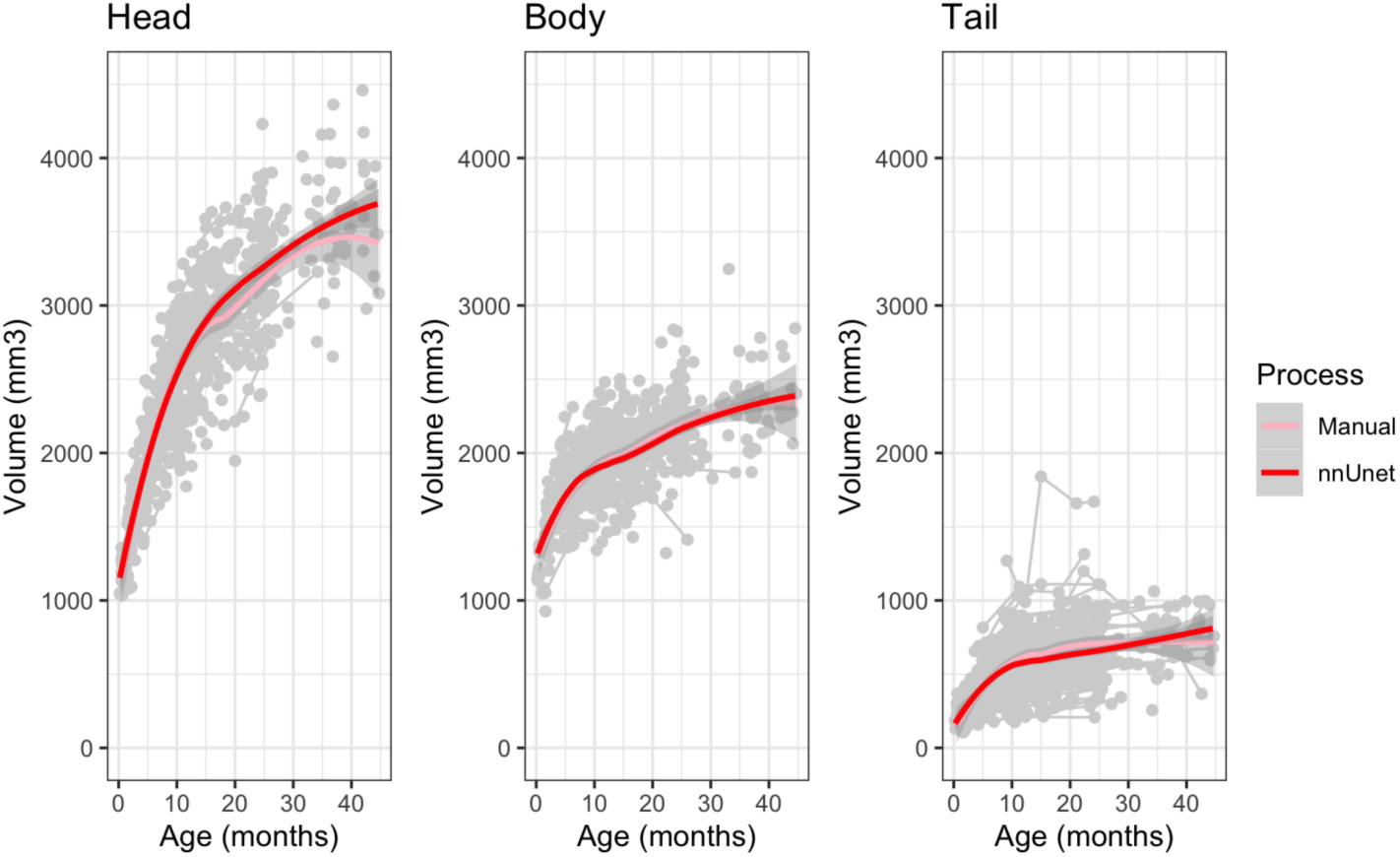
Manually and automatically (nnUnet) segmented hippocampal trajectories. Trajectories were highly concordant across all subregions for manual compared to automatic segmentations.

### Developmental Trajectories

The head of the hippocampus demonstrated hemispheric differences in trajectory (Supplemental Figure 4), so GAMMs models were created separately for the left and right hemispheres. Figure 5 shows the GAMMs models for each hippocampal subregion, demonstrating the different developmental trajectories in each. As expected, based on the first derivative of the GAMMs models, growth was fastest early in development. Significant growth was evident for longer in the head than in the body or tail of the hippocampus (Figure 5, green bars). The head of the hippocampus showed similar growth rates across left and right hemispheres, with a slightly larger volume in the right hemisphere starting early in development. To provide a quantitative measure of growth inflection, the age at which the rate of growth in the GAMMs model slowed to 50% of maximum growth was also calculated, which aligned visually with a plateau in growth. Growth leveled off in the left head by 11.5 months and by 10 months in the right head. In the body of the hippocampus, growth slowed earlier, at 5.6 months of age. The tail of the hippocampus showed similar steep growth until 7.8 months. Sex differences were seen across subregions in the expected direction, such that volumes were slightly larger in males than females, although this trend failed to reach significance in the tail (tail: p=0.090; body: p=0.034; left and right head: p<0.001).

**Figure 5.**
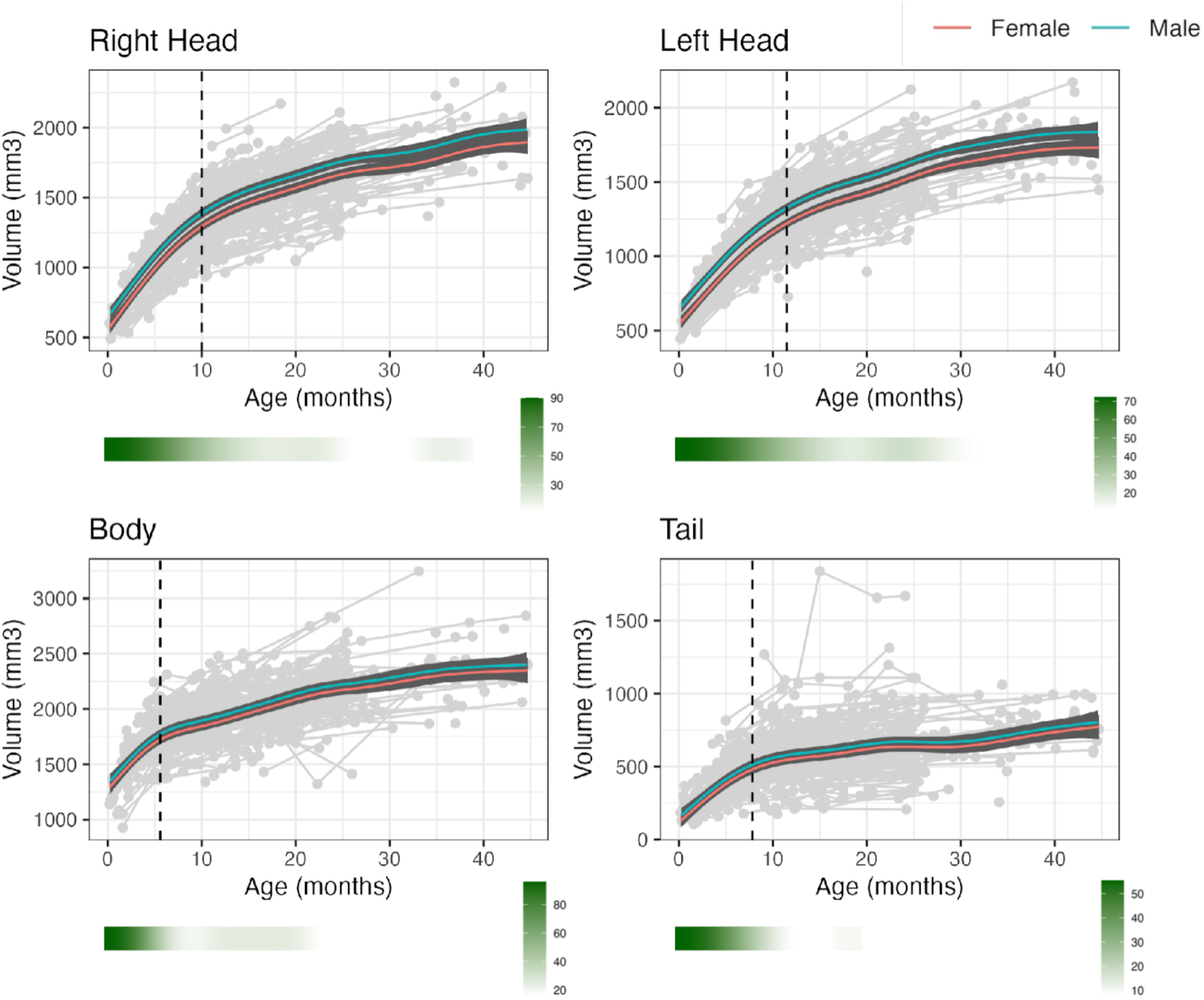
Developmental trajectories. For each region, raw data are plotted with the final GAMM model by sex, with a 95% CI. Separate models were created for the left and right hemisphere for the head of the hippocampus, which demonstrated hemispheric differences along with sex differences. The dotted line represents the age at which the rate of growth in the GAMMs model slowed to 50% of maximum growth, which varied across subregions. Green plots indicate regions of significant age-related change based on posterior simulation of model coefficients: darker green = faster growth rate.

### Sleep

BISQ sleep data was available for 902 participants in the full BCP sample (885 with total sleep duration, 901 with sleep problems), not all of whom had imaging data. In the full sleep sample, BISQ total sleep decreased from about 16 hours of sleep in the newborn period to about 12 hours of sleep by 60 months of age, as expected. The BISQ sleep problems score was consolidated from three responses, “no problem”, “small problem”, and “serious problem” to two, “no problem”, “problem”, as only 4 parents in the full sample endorsed “serious problem”. Across the sample, 19% of parents endorsed that sleep was a “problem”. In this full sample, total hours of sleep correlated with sleep problems after accounting for age and sex (F(3, 880)=75.18, p<0.001, estimate=-0.70). Among visits with BISQ sleep data as well as imaging data and later memory outcomes (N=107), the same total sleep decrease was seen over age, and a similar percentage of parents (25%) rated sleep as a problem. The relation between the two was trending (F(3, 103)=12.49, p=0.075, estimate=-0.49) in this smaller sample.

At the follow-up school age visit, all 58 participants had available sleep metrics from the CSHQ. Participants averaged 10.7 hours of sleep, which decreased across the 3–7-year-old age range.

Participants averaged a total sleep problems score of 38.8 (SD=5.7), with possible scores ranging from 31-97. There was no correlation between the sleep problems score and total sleep duration at this age.

### Source Memory

Participants on average remembered 50% of the novel facts. Of these, on average participants also remembered the source from 21% of the novel facts. Participant performance was highly variable, ranging from 0 to 58% of Fact+Source items remembered. As expected, performance correlated positively with age (F(1,52)=7.87, p=0.007). A lack of still image prompts as cues was a significant predictor of performance at floor (zero Fact+Source items correct vs. any) after adjusting for age (Logistic regression p-value for cues=0.005). Thus, the use of cues was included as a covariate in subsequent analyses. The types of source errors also changed with age. As previous studies with this task have shown (Drummey & Newcombe, 2002), the proportion of items incorrectly attributed to outside sources, rather than to guessing/just knowing or correctly attributing it to the experiment but the incorrect video, decreased with age (F(1,52)=6.48, p=0.014). After accounting for age and sex, neither total sleep nor sleep disturbance score from the CSHQ correlated with source memory performance.

### Pattern Discrimination

Figure 6A shows all participants’ performance across each of the trial types. Overall, children were very good at recognizing “same” and “new” images and worse at recognizing “similar” images. On average, children were more likely to assign a “similar” image as “same” rather than “similar”, suggesting that many of the small differences between the encoding image and test image were not remembered at this age. As seen in Figure 6B, where the participants were split into younger and older groups based on the median age, the likelihood of accurately endorsing a “similar” image increased with age, with a corresponding decrease in likelihood of inaccurately assigning the label “new” to “similar” images. Accordingly, lure discrimination index performance on the MST correlated positively with age (F(1, 55)=7.4, p<0.001). This held true after applying the recognition filter to exclude the eight participants whose performance indicated a lack of understanding or engagement with the task in its entirety.

**Figure 6.**
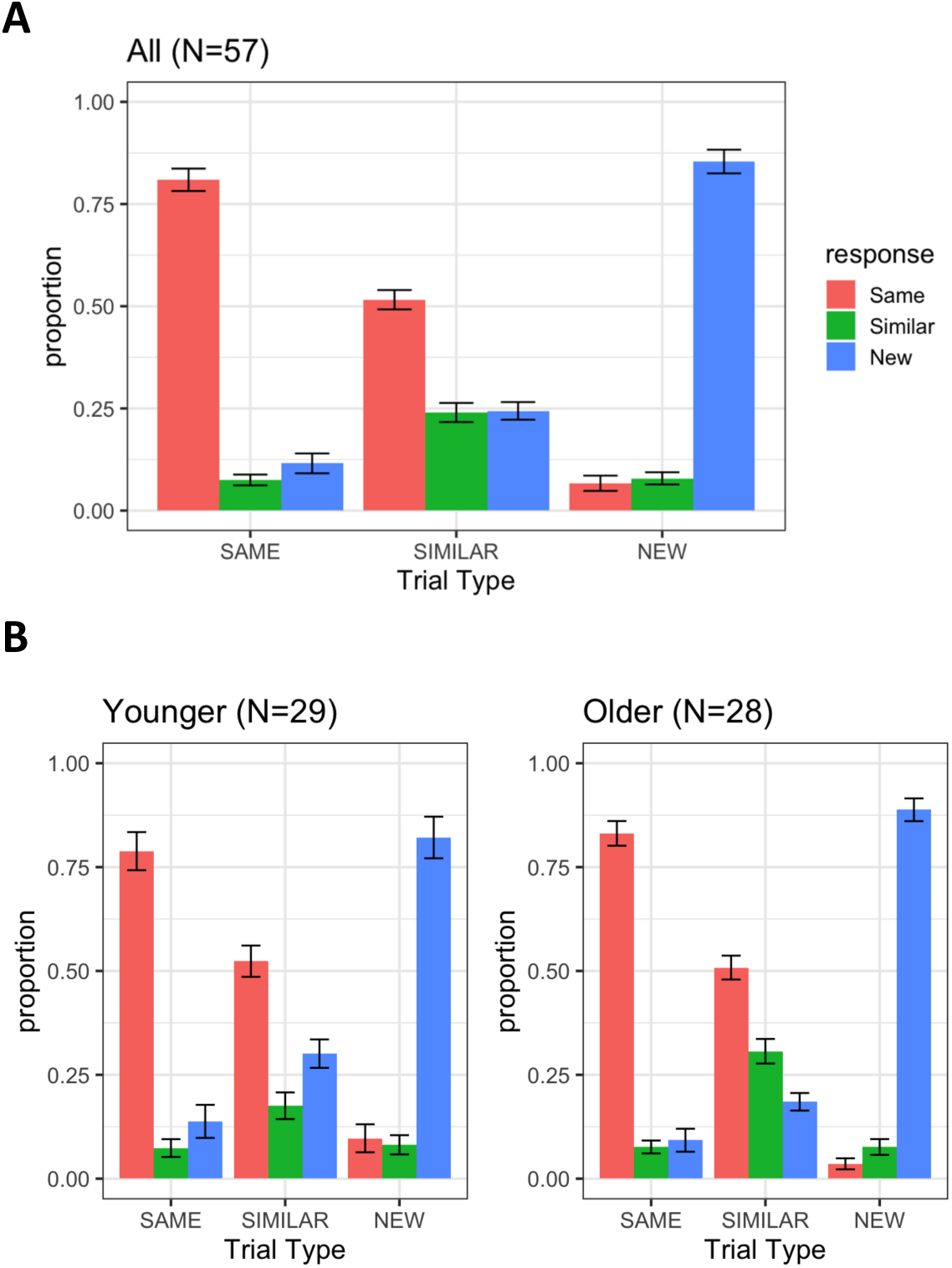
Plot of participant performance on the MST task across trial types. A) All participants, B) With a median split in age at 66.7 months to compare across age

After accounting for age and sex, total sleep from the CSHQ at the school age follow-up showed a trend correlation with performance on the MST that did not reach our threshold for significance (F(3, 53)=5.54, p=0.065). In the sensitivity analysis where the recognition filter was applied, the association dropped close to the significance threshold (F(3, 45)=10.56, p=0.053), suggesting a potential, though tenuous relationship. Sleep disturbance score from the CSHQ did not correlate with performance on the MST task.

### Brain Development Associations with Episodic Memory

On the MST, GAMMs residuals from the right head of the hippocampus showed a positive association with pattern discrimination performance (F(10, 120)=3.88, p=0.015, Table 3 top). This held after applying the recognition filter to exclude participants with poor overall recognition on the MST. When using the GAMMs residuals that included an effect of total brain volume, both the right head of the hippocampus and the right thalamus showed a significant correlation with task performance, with the head of the hippocampus positively predicting performance, and the thalamus showing a negative association (F(10, 120)=4.52, p_Rhippohead_=0.037, p_Rthalamus_=0.035, Table 4 bottom). After including the effect of FSIQ from the WPPSI-IV, only the association with the right thalamus held (F(11, 117)=4.40, p_Rhippohead_=0.111, p_Rthalamus_=0.015). The WPPSI-IV was separated into separate indices, the verbal comprehension index (comprised of information and similarities subtests), working memory index (picture memory and zoo location subtests), block design (part of the visual spatial index), and matrix reasoning (part of the fluid reasoning index). When these were added as covariates separately, the association with the right head of the hippocampus still held with each of the indices and subtests except for block design. As the block design subtest is a measure of visual spatial skills, it suggests that the FSIQ effect has more to do with the overlap in visual spatial skills between the block design task on the FSIQ and visual spatial skills also inherent in the MST task rather than a broader IQ effect.

**Table 3.** MST pattern discrimination performance models. All models include age at MRI, age at follow-up, and number of participant visits.

| Pattern Discrimination |  |  |  |  |
| --- | --- | --- | --- | --- |
| Region | estimate | SE | 95% CI | p-value |
| Left Head | 5.08e-05 | 1.87e-04 | -3.19e-04, 4.21e-04 | 0.786 |
| Right Head | 3.91e-04 | 1.58e-04 | 7.86e-05, 7.04e-04 | **0.015** |
| Body | 1.19e-04 | 1.22e-04 | -1.22e-04, 3.60e-04 | 0.331 |
| Tail | -1.59e-04 | 1.38e-04 | -4.33e-04, 1.15e-04 | 0.254 |
| Left Thalamus | 1.55e-04 | 1.43e-04 | -1.28e-04, 4.37e-04 | 0.281 |
| Right Thalamus | -2.75e-04 | 1.46e-04 | -5.65e-04, 1.48e-05 | 0.063 |
| Amygdala | 8.62e-05 | 2.30e-04 | -3.70e-04, 5.43e-04 | 0.709 |
| GAMM residuals including TBV |  |  |  |  |
| Region | estimate | SE | 95% CI | p-value |
| Left Head | 1.03e-04 | 1.79e-04 | -2.52e-04, 4.57e-04 | 0.568 |
| Right Head | 3.28e-04 | 1.55e-04 | 2.07e-05, 6.35e-04 | **0.037** |
| Body | 2.01e-04 | 1.17e-04 | -3.12e-05, 4.34e-04 | 0.089 |
| Tail | -2.39e-05 | 1.32e-04 | -2.86e-04, 2.38e-04 | 0.857 |
| Left Thalamus | 5.31e-05 | 1.39e-04 | -2.22e-04, 3.28e-04 | 0.702 |
| Right Thalamus | -3.06e-04 | 1.44e-04 | -5.90e-04, -2.16e-05 | **0.035** |
| Amygdala | -4.35e-04 | 2.32e-04 | -8.95e-04, 2.53e-05 | 0.064 |

When early sleep measures from the BISQ were added to the models, neither total sleep nor sleep problems were a significant predictor of memory performance. For school-aged sleep from the CSHQ, sleep disturbance score was not a significant predictor, but total sleep was. In that model, the right head of the hippocampus was no longer a significant predictor of MST performance, but was trending (F(11, 119)=4.39, p_Rhippohead_=0.067, p_Rthalamus_=0.047). This total model also accounted for a small amount of additional variance above and beyond the model without sleep included (®R^2^=0.04). In the model where the GAMMs were adjusted for total brain volume, only the right thalamus remained significant, with the body of the hippocampus trending and no correlation seen between the right head of the hippocampus and MST performance (F(11, 119)=5.21, p_Rhippohead_=0.144, p_body_=0.054, p_Rthalamus_=0.020). This model also accounted for a small amount of additional variance above and beyond the model without sleep included (®R^2^=0.05). Altogether, these results suggest that sleep may predict later pattern discrimination, and that some of the variance predicted may be shared by early brain development in memory networks.

No correlations were observed between early subcortical regional volumes and school-age memory performance in the source memory task.

## Discussion

Here we demonstrate distinct nonlinear longitudinal developmental trajectories across subregions of the hippocampus in the first years of life. The head of the hippocampus showed steep growth that plateaued several months later than the plateau in the tail and body. By using a framework akin to normative modelling, we determined how different a participant’s regional brain volume was from their predicted volume based on their age and sex (and total brain volume when included). This approach is consistent with many other studies of brain development (Bethlehem et al., 2022; McCormick et al., 2023), and variations of this approach have been suggested by the World Health Organization (WHO) when constructing new growth curves (Borghi et al., 2006). This sample is the most densely sampled study across these first years of life, providing important nuance to previous examinations of subcortical growth patterns, especially in memory networks. The new hippocampal subregion segmentation pipeline and the underlying manual hippocampal subregion segmentations are available to the scientific community as an open resource to automatically segment hippocampal subregions in infants across other studies of early hippocampal development.

Differences in hippocampal definitions make comparison across studies difficult. Overall hippocampal volumes from this sample are consistent with some prior studies (Canada et al., 2020; Gilmore et al., 2012), but showed qualitative differences in volume or trajectory from other prior datasets (Chen et al., 2023; Choe et al., 2013; Nichols, Grace, et al., 2023; Uematsu et al., 2012). These discrepancies may depend both on differences in hippocampal definitions (e.g. inclusion of the full length of the hippocampal tail or not) and the density with which participants were sampled in the first years of life, which was limited in many of lifespan studies (Uematsu et al., 2012). Studies using the same protocol for the definition of the hippocampus show similar volumes for the youngest of their participants at 4 years of age (Canada et al., 2020) compared to the oldest participants in our sample (who are just under 4 years old), including across the head, body, and tail subregions. Work is underway to standardize the definition of hippocampal structures (Canada et al., 2024; Wolf et al., 2017), but these efforts are focused primarily on adult and aging populations currently. Such work needs to be expanded to the youngest populations as well, who present different challenges for imaging, including increased motion, smaller structures, and changing image contrasts due to myelination processes. Hemispheric differences observed in the volume of the hippocampal head are consistent with previous studies that have shown slightly larger right than left hippocampi (Holland et al., 2014; Pfluger et al., 1999; Thompson et al., 2009), but this laterality is not consistent across the literature, e.g. (Gilmore et al., 2012). Moving forward, estimated hippocampal trajectories should be confirmed using early life datasets beyond the BCP, such as the Infant Brain Imaging Study (IBIS) dataset (Burrows et al., 2024) and the ongoing Healthy Brain and Child Development (HBCD) dataset (Dean et al., 2024).

Specific timing of growth trajectory changes, such as the growth rate changes observed near 6, 8, and 10-12 months of age across the hippocampal subregions, may provide new insight into the timing and mechanisms of hippocampal dependent episodic memory development in the first years of life. For example, the changes in timing of hippocampal development align closely with behavioral findings on episodic memory. Nine months of age is sometimes considered a transition period for episodic memory, when memory can begin to be measured with tasks that explicitly tap episodic memory (Kagan & Hamburg, 1981; Mullally & Maguire, 2014; Richmond & Nelson, 2009). It could be that a minimum baseline of growth may be needed before clear episodic memory can be demonstrated, including dendritic and synaptic development, axonal growth, and glial proliferation processes that are collapsed into structural volumes metrics.

Additionally, we found that the right head of the hippocampus and the right thalamus predicted early school-age pattern discrimination ability. Positive correlations between hippocampal volumes and episodic memory are consistent with meta-analyses showing that larger overall hippocampal volumes are correlated with better memory performance in children and adolescents (Botdorf et al., 2022). The brain-behavior associations found in our study differed by region/subregion (e.g. hippocampus vs. amygdala and head vs. tail of the hippocampus), hemisphere (right vs. left hemisphere), and type of memory (pattern discrimination vs. source memory). These differences suggest that relations between brain growth and memory are region and context specific, as has been previously demonstrated in older children. In children, volume in the CA2-4/DG subfields has been reported to correlate with pattern discrimination performance, while CA1 and CA2-4/DG in the head and body have been shown to correlate with source memory performance (Canada et al., 2019, 2021; Riggins et al., 2015, 2018). These associations differ in magnitude and directionality across age, for example, in some cases, showing positive associations in younger children and negative associations in older children. Given that our analyses are in an even younger population, positive hippocampal associations are consistent with these prior studies. Yet prior studies have also found bilateral hippocampal-memory relationships as well as hippocampal relationships with source memory performance, which we did not find in these analyses. Subregion definitions of head, body, and tail are sometimes used as proxies for subfields, given the disproportionate distribution of subfields across the longitudinal axis, with more of the CA fields in the head and more of the dentate gyrus in the body/tail (Poppenk et al., 2013). However, this is a gross proxy for subfields, especially given that studies with sufficiently high resolution to parse subfields separately within each of the subregions find differences in function and structure across both subfields and subregions in adults, with separate functions for anterior CA fields vs. posterior CA fields, for example (Beer et al., 2018; Travis et al., 2014). Studies in adults have shown that smaller anterior and larger posterior volumes of the hippocampus correlate with better performance across episodic memory tasks (Poppenk & Moscovitch, 2011), but again, alternative associations have been found in children and adolescents. In our findings, we 1) cannot parse hippocampal subfields, 2) are predicting later rather than concurrent memory, and 3) are studying these relations earlier in development than any of the studies cited above, so it is unsurprising that our results show a similar but slightly different pattern of results than prior literature. Differential behavioral correlations during early development might be because the hippocampus is supporting different developmental processes occurring at that time, such that specific regions that are important for episodic memory during adulthood might be different than those important during infancy or childhood. While more work replicating these findings is needed to determine the stability and reproducibility of these brain-behavior associations in early development, these results provide insight towards how protracted brain development may relate to the development of episodic memory abilities in infants and young children.

The right-sided laterality of the hippocampal and thalamic associations with later episodic memory in our study is inconsistent with previous literature in adults. In adults, literature suggests that episodic memory is more strongly associated with the left hippocampus, while the right is more associated with spatial memory (Burgess et al., 2002). However, this laterality difference depends on the specific memory task, the temporal remoteness of the memory, and the age of the participant, so it’s possible that the right hippocampus plays a larger role earlier in development, and becomes more left lateralized by young adulthood (Hopf et al., 2013; Maguire & Frith, 2003). Additionally, episodic memory tasks used in adults are often verbal, so it’s possible that lateralization in the adult literature is picking up on verbal/semantic lateralization to the left hemisphere, while the MST is more object-based and thus right medial-temporal lobe dependent.

Findings in the thalamus suggest that a smaller thalamus early in development predicts better pattern discrimination at school age. This negative association with the thalamus is potentially unexpected at first glance, but could indicate a trade-off between nearby structural regions, including the hippocampus, especially since significant associations with the thalamus were seen only after accounting for total brain volume. More work is needed to understand the role of the thalamus in early episodic memory development. No associations were found between early amygdala development and later episodic memory. Considering that the amygdala is most strongly associated with processing the emotional valence of memories, it’s not surprising that we did not see anything with the relatively neutral, unemotional stimuli used in our two episodic memory tasks.

No correlations were found between early brain development and performance on the source memory task. It’s possible this represents a true null finding. The source memory task measures different aspects of episodic memory than the MST task, including focusing on source context vs. pattern discrimination, and focusing on longer-term memory retrieval (week delay) than the MST (delay of just a few minutes). As well, as the analyses with behavioral outcomes were performed on a subset of participants with follow-up data (N=58), these analyses are less powered than the overall trajectory analyses. Alternatively, the source memory task was confounded by the addition of cues partway into data collection. This was necessary considering the floor performance of many of participants before the implementation of cues, but did create an additional source of variation that might have made it harder to detect small brain-behavior relations. The age range we measured, 0-5 years, is quite a wide range and reflects a dynamic period in development. Therefore, it is also possible that unique predictive associations at different ages were lost once ages were combined. Future analyses could attempt to examine separate age groups that might demonstrate different predictive associations with later episodic memory performance, potentially surrounding the inflection points of growth identified in the current dataset.

Total sleep duration at the early school age visit accounted for variance in pattern discrimination performance above and beyond brain volume correlations. The hippocampal correlations, especially in the right head, were not robust to the addition of this sleep measure. Thus, it’s possible these hippocampal-memory and sleep-memory correlations are pulling from similar pools of variance. Future analyses should look more formally at the potential mediating or moderating associations between sleep, brain development, and memory to disentangle any overlapping associations.

This study focused on structural development, but an important next step would be to analyze patterns of connectivity, which conceptually may be more closely tied to later behavior than structural volumes. Work in adults has shown differential connectivity patterns to the default mode network (DMN) vs. the parietal memory network (PMN) across the longitudinal axis of the hippocampus, with anterior portions more closely connected to the DMN, and posterior portions more strongly connected to the PMN (Zheng et al., 2021). A recent paper suggested that long-axis connectivity specialization may be seen even as early as the fetal period (Nichols et al., 2025). An understanding of the developmental roots of this specialization along the hippocampus will be important to understand memory network development.

As this study used early brain development to predict later memory performance, these results cannot address how brain development relates to memory performance at the time of the change in growth. Future studies should aim to measure concurrent associations between brain region volumes and memory across the first years of life. This could provide more insight into how the timing of hippocampal growth relates to more immediate changes in episodic memory performance across the ages involved in the infantile/childhood amnesia phenomenon. Ages of plateauing growth found in this study could provide hypothesized ages for close analysis in a future study of these concurrent associations.

The identified associations between brain development and memory function in typically developing children suggest opportunities to use brain imaging methods to identify potential predictors of episodic memory performance in other populations at risk for altered brain and behavioral development. Discovering how and when these memory networks can be disrupted by early brain insults may provide a deeper understanding of the neural mechanisms of typical early memory development and provide opportunities to improve clinical care in such cases. Further research is needed on the causal associations between brain and behavioral development following neonatal insults, which first necessitates a more complete understanding of typical trajectories of development. Together, normative and altered developmental trajectories may inform new treatments and interventions for neonates with early insults to these developing systems.

## Acknowledgements

Baby Connectome Project data collection was supported by grants R01-MH104324 and U01-MH110274 and the Bill & Melinda Gates Foundation INV-015711. SMS was supported by the National Science Foundation Graduate Research Fellowship Program (2237827). Portions of this manuscript contributed to SMS’s doctoral dissertation. We would like to thank Dr. Tracy Riggins for the use of her image presentation materials for the Source Memory and Mnemonic Similarities Tasks.

## Code Accessibility

Code related to this manuscript, including links to the trained hippocampal segmentation model, will be available at https://github.com/sallystoyell/Hippocampal_Subregion_Development.

## Data Availability

BCP data is available through the NDA (https://nda.nih.gov/edit_collection.html?id=2848). Manual and automatic hippocampal subregion segmentations will be available at the MIDB Open Repository.

## Author Contributions

SMS: Designed research, performed research, analyzed data, wrote the first draft of the paper, wrote the paper, edited the paper

JTL: Analyzed data

LH: Performed research

AB: Performed research

AB: Performed research

KMT: Edited the paper

DAF: Edited the paper

BT-C: Analyzed data, edited the paper

EF: Designed research, analyzed data, edited the paper

JTE: Designed research, edited the paper

## References

Beer, Z., Vavra, P., Atucha, E., Rentzing, K., Heinze, H.-J., & Sauvage, M. M. (2018). The memory for time and space differentially engages the proximal and distal parts of the hippocampal subfields CA1 and CA3. PLOS Biology, 16(8), e2006100. 10.1371/journal.pbio.2006100

Bethlehem, R. A. I., Seidlitz, J., White, S. R., Vogel, J. W., Anderson, K. M., Adamson, C., Adler, S., Alexopoulos, G. S., Anagnostou, E., Areces-Gonzalez, A., Astle, D. E., Auyeung, B., Ayub, M., Bae, J., Ball, G., Baron-Cohen, S., Beare, R., Bedford, S. A., Benegal, V., … Alexander-Bloch, A. F. (2022). Brain charts for the human lifespan. Nature, 604(7906), 525–533. 10.1038/s41586-022-04554-y

Bevandić, J., Chareyron, L. J., Bachevalier, J., Cacucci, F., Genzel, L., Newcombe, N. S., Vargha-Khadem, F., & Ólafsdóttir, H. F. (2024). Episodic memory development: Bridging animal and human research. Neuron, 112(7), 1060–1080. 10.1016/j.neuron.2024.01.020

Billot, B., Greve, D. N., Puonti, O., Thielscher, A., Van Leemput, K., Fischl, B., Dalca, A. V., & Iglesias, J. E. (2023). SynthSeg: Segmentation of brain MRI scans of any contrast and resolution without retraining. Medical Image Analysis, 86, 102789. 10.1016/j.media.2023.102789

Borghi, E., de Onis, M., Garza, C., Van den Broeck, J., Frongillo, E. A., Grummer-Strawn, L., Van Buuren, S., Pan, H., Molinari, L., Martorell, R., Onyango, A. W., Martines, J. C., & Group, for the W. M. G. R. S. (2006). Construction of the World Health Organization child growth standards: Selection of methods for attained growth curves. Statistics in Medicine, 25(2), 247–265. 10.1002/sim.2227

Botdorf, M., Canada, K. L., & Riggins, T. (2022). A meta-analysis of the relation between hippocampal volume and memory ability in typically developing children and adolescents. Hippocampus, 32(5), 386–400. 10.1002/hipo.23414

Bouyeure, A., Patil, S., Mauconduit, F., Poiret, C., Isai, D., & Noulhiane, M. (2021). Hippocampal subfield volumes and memory discrimination in the developing brain. Hippocampus, 31(11), 1202–1214. 10.1002/hipo.23385

Burgess, N., Maguire, E. A., & O’Keefe, J. (2002). The human hippocampus and spatial and episodic memory. Neuron, 35(4), 625–641. 10.1016/s0896-6273(02)00830-9

Burrows, C. A., Lasch, C., Gross, J., Girault, J. B., Rutsohn, J., Wolff, J. J., Swanson, M. R., Lee, C. M., Dager, S. R., Cornea, E., Stephens, R., Styner, M., John, T. St., Pandey, J., Deva, M., Botteron, K. N., Estes, A. M., Hazlett, H. C., Pruett, J. R., … Elison, J. T. (2024). Associations between early trajectories of amygdala development and later school-age anxiety in two longitudinal samples. Developmental Cognitive Neuroscience, 65, 101333. 10.1016/j.dcn.2023.101333

Callow, D. D., Canada, K. L., & Riggins, T. (2020). Microstructural Integrity of the Hippocampus During Childhood: Relations With Age and Source Memory. Frontiers in Psychology, 11, 568953. 10.3389/fpsyg.2020.568953

Canada, K. L., Botdorf, M., & Riggins, T. (2020). Longitudinal development of hippocampal subregions from early- to mid-childhood. Hippocampus, 30(10), 1098–1111. 10.1002/hipo.23218

Canada, K. L., Hancock, G. R., & Riggins, T. (2021). Modeling longitudinal changes in hippocampal subfields and relations with memory from early- to mid-childhood. Developmental Cognitive Neuroscience, 48, 100947. 10.1016/j.dcn.2021.100947

Canada, K. L., Mazloum-Farzaghi, N., Rådman, G., Adams, J. N., Bakker, A., Baumeister, H., Berron, D., Bocchetta, M., Carr, V. A., Dalton, M. A., de Flores, R., Keresztes, A., La Joie, R., Mueller, S. G., Raz, N., Santini, T., Shaw, T., Stark, C. E. L., Tran, T. T., … Group, the H. S. (2024). A (sub)field guide to quality control in hippocampal subfield segmentation on high-resolution T-weighted MRI. Human Brain Mapping, 45(15), e70004. 10.1002/hbm.70004

Canada, K. L., Ngo, C. T., Newcombe, N. S., Geng, F., & Riggins, T. (2019). It’s All in the Details: Relations Between Young Children’s Developing Pattern Separation Abilities and Hippocampal Subfield Volumes. Cerebral Cortex, 29(8), 3427–3433. 10.1093/CERCOR/BHY211

Chen, L., Wang, Y., Wu, Z., Shan, Y., Li, T., Hung, S.-C., Xing, L., Zhu, H., Wang, L., Lin, W., & Li, G. (2023). Four-dimensional mapping of dynamic longitudinal brain subcortical development and early learning functions in infants. Nature Communications, 14(1), 3727. 10.1038/s41467-023-38974-9

Choe, M. S., Ortiz-Mantilla, S., Makris, N., Gregas, M., Bacic, J., Haehn, D., Kennedy, D., Pienaar, R., Caviness, V. S., Benasich, A. A., & Ellen Grant, P. (2013). Regional infant brain development: An MRI-based morphometric analysis in 3 to 13 month olds. Cerebral Cortex, 23(9), 2100–2117. 10.1093/cercor/bhs197

Dean, D. C., Tisdall, M. D., Wisnowski, J. L., Feczko, E., Gagoski, B., Alexander, A. L., Edden, R. A. E., Gao, W., Hendrickson, T. J., Howell, B. R., Huang, H., Humphreys, K. L., Riggins, T., Sylvester, C. M., Weldon, K. B., Yacoub, E., Ahtam, B., Beck, N., Banerjee, S., … Elison, J. T. (2024). Quantifying brain development in the HEALthy Brain and Child Development (HBCD) Study: The magnetic resonance imaging and spectroscopy protocol. Developmental Cognitive Neuroscience, 70, 101452. 10.1016/j.dcn.2024.101452

Decker, A. L., Duncan, K., Finn, A. S., & Mabbott, D. J. (2020). Children’s family income is associated with cognitive function and volume of anterior not posterior hippocampus. Nature Communications, 11(1). 10.1038/s41467-020-17854-6

DeMaster, D., Pathman, T., Lee, J. K., & Ghetti, S. (2014). Structural Development of the Hippocampus and Episodic Memory: Developmental Differences Along the Anterior/Posterior Axis. Cerebral Cortex, 24(11), 3036–3045. 10.1093/CERCOR/BHT160

Dick, A. S., Ralph, Y., Farrant, K., Reeb-Sutherland, B., Pruden, S., Mattfeld, A. T., & Dick, S. (2022). Volumetric development of hippocampal subfields and hippocampal white matter connectivity: Relationship with episodic memory. Developmental Psychobiology, 64(8), e22333–e22333. 10.1002/DEV.22333

Drummey, A. B., & Newcombe, N. S. (2002). Developmental changes in source memory. Developmental Science, 5(4), 502–513. 10.1111/1467-7687.00243

Ellis, C. T., Skalaban, L. J., Yates, T. S., Bejjanki, V. R., Córdova, N. I., & Turk-Browne, N. B. (2021). Evidence of hippocampal learning in human infants. Current Biology, 31(15), 3358–3364.e4. 10.1016/j.cub.2021.04.072

Feczko, E., Stoyell, S. M., Moore, L. A., Alexopoulos, D., Bagonis, M., Barrett, K., Bower, B., Cavender, A., Chamberlain, T. A., Conan, G., Day, T. K. M., Goradia, D., Graham, A., Heisler-Roman, L., Hendrickson, T. J., Houghton, A., Kardan, O., Kiffmeyer, E. A., Lee, E. G., … Elison, J. T. (2025). Baby Open Brains: An open-source dataset of infant brain segmentations. Scientific Data, 12, 1423. 10.1038/s41597-025-05404-y

Ghetti, S., & Bunge, S. A. (2012). Neural changes underlying the development of episodic memory during middle childhood. Developmental Cognitive Neuroscience, 2(4), 381–395. 10.1016/j.dcn.2012.05.002

Gilmore, J. H., Shi, F., Woolson, S. L., Knickmeyer, R. C., Short, S. J., Lin, W., Zhu, H., Hamer, R. M., Styner, M., & Shen, D. (2012). Longitudinal development of cortical and subcortical gray matter from birth to 2 years. Cerebral Cortex, 22(11), 2478–2485. 10.1093/cercor/bhr327

Gómez, R. L., & Edgin, J. O. (2016). The extended trajectory of hippocampal development: Implications for early memory development and disorder. Developmental Cognitive Neuroscience, 18, 57–69. 10.1016/j.dcn.2015.08.009

Hendrickson, T. J., Reiners, P., Moore, L. A., Lundquist, J. T., Fayzullobekova, B., Perrone, A. J., Lee, E. G., Moser, J., Day, T. K. M., Alexopoulos, D., Styner, M., Kardan, O., Chamberlain, T. A., Mummaneni, A., Caldas, H. A., Bower, B., Stoyell, S., Martin, T., Sung, S., … Feczko, E. (2025). BIBSNet: A Deep Learning Baby Image Brain Segmentation Network for MRI Scans. bioRxiv, 2023.03.22.533696. 10.1101/2023.03.22.533696

Holland, D., Chang, L., Ernst, T. M., Curran, M., Buchthal, S. D., Alicata, D., Skranes, J., Johansen, H., Hernandez, A., Yamakawa, R., Kuperman, J. M., & Dale, A. M. (2014). Structural Growth Trajectories and Rates of Change in the First 3 Months of Infant Brain Development. JAMA Neurology, 71(10), 1266–1274. 10.1001/JAMANEUROL.2014.1638

Hopf, L., Quraan, M. A., Cheung, M. J., Taylor, M. J., Ryan, J. D., & Moses, S. N. (2013). Hippocampal Lateralization and Memory in Children and Adults. Journal of the International Neuropsychological Society, 19(10), 1042–1052. 10.1017/S1355617713000751

Howell, B. R., Styner, M. A., Gao, W., Yap, P. T., Wang, L., Baluyot, K., Yacoub, E., Chen, G., Potts, T., Salzwedel, A., Li, G., Gilmore, J. H., Piven, J., Smith, J. K., Shen, D., Ugurbil, K., Zhu, H., Lin, W., & Elison, J. T. (2019). The UNC/UMN Baby Connectome Project (BCP): An overview of the study design and protocol development. NeuroImage, 185, 891–905. 10.1016/j.neuroimage.2018.03.049

Huber, R., & Born, J. (2014). Sleep, synaptic connectivity, and hippocampal memory during early development. Trends in Cognitive Sciences, 18(3), 141–152. 10.1016/j.tics.2013.12.005

Isensee, F., Jaeger, P. F., Kohl, S. A. A., Petersen, J., & Maier-Hein, K. H. (2021). nnU-Net: A self-configuring method for deep learning-based biomedical image segmentation. Nature Methods, 18(2), 203–211. 10.1038/s41592-020-01008-z

Jabès, A., & Nelson, C. A. (2015). 20 years after “The ontogeny of human memory: A cognitive neuroscience perspective,” where are we? International Journal of Behavioral Development, 39(4), 293–303. 10.1177/0165025415575766

Johnson, E. G., Mooney, L., Graf Estes, K., Nordahl, C. W., & Ghetti, S. (2021). Activation for newly learned words in left medial-temporal lobe during toddlers’ sleep is associated with memory for words. Current Biology, 31(24), 5429–5438.e5. 10.1016/j.cub.2021.09.058

Kagan, J., & Hamburg, M. (1981). The Enhancement of Memory in the First Year. The Journal of Genetic Psychology, 138(1), 3–14. 10.1080/00221325.1981.10532837

Kurdziel, L., Duclos, K., & Spencer, R. M. C. (2013). Sleep spindles in midday naps enhance learning in preschool children. Proceedings of the National Academy of Sciences of the United States of America, 110(43), 17267–17272. 10.1073/pnas.1306418110

Langnes, E., Sneve, M. H., Sederevicius, D., Amlien, I. K., Walhovd, K. B., & Fjell, A. M. (2020). Anterior and posterior hippocampus macro- and microstructure across the lifespan in relation to memory—A longitudinal study. Hippocampus, 30(7), 678–692. 10.1002/hipo.23189

Lavenex, P., & Banta Lavenex, P. (2013). Building hippocampal circuits to learn and remember: Insights into the development of human memory. Behavioural Brain Research, 254, 8–21. 10.1016/j.bbr.2013.02.007

Lee, J. K., Fandakova, Y., Johnson, E. G., Cohen, N. J., Bunge, S. A., & Ghetti, S. (2020). Changes in anterior and posterior hippocampus differentially predict item-space, item-time, and item-item memory improvement. Developmental Cognitive Neuroscience, 41, 100741–100741. 10.1016/J.DCN.2019.100741

Li, G., Chen, M.-H., Li, G., Wu, D., Lian, C., Sun, Q., Rushmore, R. J., & Wang, L. (2023). Volumetric Analysis of Amygdala and Hippocampal Subfields for Infants with Autism. Journal of Autism and Developmental Disorders, 53(6), 2475–2489. 10.1007/s10803-022-05535-w

Lokhandwala, S., & Spencer, R. M. C. (2021). Slow wave sleep in naps supports episodic memories in early childhood. Developmental Science, 24(2). 10.1111/desc.13035

Luby, J. L., Barch, D. M., Belden, A., Gaffrey, M. S., Tillman, R., Babb, C., Nishino, T., Suzuki, H., & Botteron, K. N. (2012). Maternal support in early childhood predicts larger hippocampal volumes at school age. Proceedings of the National Academy of Sciences, 109(8), 2854–2859. 10.1073/pnas.1118003109

MacDuffie, K. E., Shen, M. D., Dager, S. R., Styner, M. A., Kim, S. H., Paterson, S., Pandey, J., St John, T., Elison, J. T., Wolff, J. J., Swanson, M. R., Botteron, K. N., Zwaigenbaum, L., Piven, J., & Estes, A. M. (2020). Sleep onset problems and subcortical development in infants later diagnosed with autism spectrum disorder. American Journal of Psychiatry, 177(6), 518–525. 10.1176/appi.ajp.2019.19060666

Maguire, E. A., & Frith, C. D. (2003). Lateral Asymmetry in the Hippocampal Response to the Remoteness of Autobiographical Memories. The Journal of Neuroscience, 23(12), 5302–5307. 10.1523/JNEUROSCI.23-12-05302.2003

McCormick, E. M., Byrne, M. L., Flournoy, J. C., Mills, K. L., & Pfeifer, J. H. (2023). The Hitchhiker’s guide to longitudinal models: A primer on model selection for repeated-measures methods. Developmental Cognitive Neuroscience, 63, 101281. 10.1016/j.dcn.2023.101281

Mooney, L. N., Johnson, E. G., Prabhakar, J., & Ghetti, S. (2021). Memory-related hippocampal activation during sleep and temporal memory in toddlers. Developmental Cognitive Neuroscience, 47, 100908–100908. 10.1016/J.DCN.2020.100908

Mullally, S. L., & Maguire, E. A. (2014). Learning to remember: The early ontogeny of episodic memory. Developmental Cognitive Neuroscience, 9, 12–29. 10.1016/j.dcn.2013.12.006

Ngo, C. T., Newcombe, N. S., & Olson, I. R. (2018). The ontogeny of relational memory and pattern separation. Developmental Science, 21(2), e12556–e12556. 10.1111/DESC.12556

Nichols, E. S., Al-Saoud, S., Fang, M., Eagleson, R., de Vrijer, B., McKenzie, C., de Ribaupierre, S., & Duerden, E. G. (2025). Early functional organization of the anterior and posterior hippocampus in the fetal brain. Cerebral Cortex (New York, NY), 35(12), bhaf327. 10.1093/cercor/bhaf327

Nichols, E. S., Blumenthal, A., Kuenzel, E., Skinner, J. K., & Duerden, E. G. (2023). Hippocampus long-axis specialization throughout development: A meta-analysis. Human Brain Mapping, 44(11), 4211–4224. 10.1002/hbm.26340

Nichols, E. S., Grace, M., Correa, S., de Vrijer, B., Eagleson, R., McKenzie, C. A., de Ribaupierre, S., & Duerden, E. G. (2023). Sex- and age-based differences in fetal and early childhood hippocampus maturation: A cross-sectional and longitudinal analysis. *Cerebral Cortex (New York*, NY), 34(1), bhad421. 10.1093/cercor/bhad421

Owens, J. A., Spirito, A., & McGuinn, M. (2000). The Children’s Sleep Habits Questionnaire (CSHQ): Psychometric properties of a survey instrument for school-aged children. Sleep, 23(8), 1043–1051. 10.1093/sleep/23.8.1d

Pfluger, T., Weil, S., Weis, S., Vollmar, C., Heiss, D., Egger, J., Scheck, R., & Hahn, K. (1999). Normative Volumetric Data of the Developing Hippocampus in Children Based on Magnetic Resonance Imaging. Epilepsia, 40(4), 414–423. 10.1111/j.1528-1157.1999.tb00735.x

Poppenk, J., Evensmoen, H. R., Moscovitch, M., & Nadel, L. (2013). Long-axis specialization of the human hippocampus. Trends in Cognitive Sciences, 17(5), 230–240. 10.1016/j.tics.2013.03.005

Poppenk, J., & Moscovitch, M. (2011). A Hippocampal Marker of Recollection Memory Ability among Healthy Young Adults: Contributions of Posterior and Anterior Segments. Neuron, 72(6), 931–937. 10.1016/j.neuron.2011.10.014

Prabhakar, J., Johnson, E. G., Nordahl, C. W., & Ghetti, S. (2018). Memory-related hippocampal activation in the sleeping toddler. Proceedings of the National Academy of Sciences of the United States of America, 115(25), 6500–6505. 10.1073/pnas.1805572115

Richmond, J., & Nelson, C. A. (2009). Relational memory during infancy: Evidence from eye tracking. Developmental Science, 12(4), 549–556. 10.1111/j.1467-7687.2009.00795.x

Riggins, T., Blankenship, S. L., Mulligan, E., Rice, K., & Redcay, E. (2015). Developmental Differences in Relations Between Episodic Memory and Hippocampal Subregion Volume During Early Childhood. Child Development, 86(6), 1710–1718. 10.1111/cdev.12445

Riggins, T., Geng, F., Botdorf, M., Canada, K., Cox, L., & Hancock, G. R. (2018). Protracted hippocampal development is associated with age-related improvements in memory during early childhood. NeuroImage, 174, 127–137. 10.1016/j.neuroimage.2018.03.009

Riggins, T., & Spencer, R. M. C. (2020). Habitual sleep is associated with both source memory and hippocampal subfield volume during early childhood. Scientific Reports, 10(1), 1–9. 10.1038/s41598-020-72231-z

Roeske, M. J., Konradi, C., Heckers, S., & Lewis, A. S. (2021). Hippocampal volume and hippocampal neuron density, number and size in schizophrenia: A systematic review and meta-analysis of postmortem studies. Molecular Psychiatry, 26(7), 3524–3535. 10.1038/s41380-020-0853-y

Sadeh, A. (2004). A brief screening questionnaire for infant sleep problems: Validation and findings for an Internet sample. Pediatrics, 113(6). 10.1542/peds.113.6.e570

Schlichting, M. L., Guarino, K. F., Schapiro, A. C., Turk-Browne, N. B., & Preston, A. R. (2017). Hippocampal Structure Predicts Statistical Learning and Associative Inference Abilities during Development. Journal of Cognitive Neuroscience, 29(1), 37–51. 10.1162/jocn_a_01028

Seehagen, S., Konrad, C., Herbert, J. S., & Schneider, S. (2015). Timely sleep facilitates declarative memory consolidation in infants. Proceedings of the National Academy of Sciences of the United States of America, 112(5), 1625–1629. 10.1073/pnas.1414000112

Simpson, G. L. (2024). gratia: An R package for exploring generalized additive models. Journal of Open Source Software, 9(104), 6962. 10.21105/joss.06962

Stark, S. M., Kirwan, C. B., & Stark, C. E. L. (2019). Mnemonic Similarity Task: A Tool for Assessing Hippocampal Integrity. Trends in Cognitive Sciences, 23(11), 938–951. 10.1016/J.TICS.2019.08.003

Tamnes, C. K., Walhovd, K. B., Engvig, A., Grydeland, H., Krogsrud, S. K., Østby, Y., Holland, D., Dale, A. M., & Fjell, A. M. (2014). Regional Hippocampal Volumes and Development Predict Learning and Memory. Developmental Neuroscience, 36(3–4), 161–174. 10.1159/000362445

Tervo-Clemmens, B., Calabro, F. J., Parr, A. C., Fedor, J., Foran, W., & Luna, B. (2023). A canonical trajectory of executive function maturation from adolescence to adulthood. Nature Communications, 14(1), 6922. 10.1038/s41467-023-42540-8

Thompson, D. K., Wood, S. J., Doyle, L. W., Warfield, S. K., Egan, G. F., & Inder, T. E. (2009). MR-determined hippocampal asymmetry in full-term and preterm neonates. Hippocampus, 19(2), 118–123. 10.1002/hipo.20492

Travis, S. G., Huang, Y., Fujiwara, E., Radomski, A., Olsen, F., Carter, R., Seres, P., & Malykhin, N. V. (2014). High field structural MRI reveals specific episodic memory correlates in the subfields of the hippocampus. Neuropsychologia, 53, 233–245. 10.1016/j.neuropsychologia.2013.11.016

Uematsu, A., Matsui, M., Tanaka, C., Takahashi, T., Noguchi, K., Suzuki, M., & Nishijo, H. (2012). Developmental Trajectories of Amygdala and Hippocampus from Infancy to Early Adulthood in Healthy Individuals. PLoS ONE, 7(10), e46970–e46970. 10.1371/journal.pone.0046970

Utsunomiya, H., Takano, K., Okazaki, M., & Mitsudome, A. (1999). Development of the temporal lobe in infants and children: Analysis by MR-based volumetry. American Journal of Neuroradiology, 20(4), 717–723.

van den Berg, N. H., Benoit, A., Toor, B., & Fogel, S. (2019). Chapter 30—Sleep Stages and Neural Oscillations: A Window into Sleep’s Role in Memory Consolidation and Cognitive Abilities. In H. C. B. T.-H. of B. N. Dringenberg (Ed.), Handbook of Sleep Research (Vol. 30, pp. 455–470). Elsevier. 10.1016/B978-0-12-813743-7.00030-X

Vijayakumar, N., Mills, K. L., Alexander-Bloch, A., Tamnes, C. K., & Whittle, S. (2018). Structural brain development: A review of methodological approaches and best practices. Developmental Cognitive Neuroscience, 33, 129–148. 10.1016/j.dcn.2017.11.008

Vinci-Booher, S., Schlichting, M. L., Preston, A. R., & Pestilli, F. (2023). Development of human hippocampal subfield microstructure and relation to associative inference. *Cerebral Cortex (New York*, NY), 33(18), 10207–10220. 10.1093/cercor/bhad276

Wang, Y., Haghpanah, F. S., Zhang, X., Santamaria, K., da Costa Aguiar Alves, G. K., Bruno, E., Aw, N., Maddocks, A., Duarte, C. S., Monk, C., Laine, A., Posner, J., & program collaborators for Environmental influences on Child Health Outcomes. (2022). ID-Seg: An infant deep learning-based segmentation framework to improve limbic structure estimates. Brain Informatics, 9(1), 12. 10.1186/s40708-022-00161-9

Wolf, D., Bocchetta, M., Preboske, G. M., Boccardi, M., Grothe, M. J., & Alzheimer’s Disease Neuroimaging Initiative. (2017). Reference standard space hippocampus labels according to the European Alzheimer’s Disease Consortium-Alzheimer’s Disease Neuroimaging Initiative harmonized protocol: Utility in automated volumetry. Alzheimer’s & Dementia: The Journal of the Alzheimer’s Association, 13(8), 893–902. 10.1016/j.jalz.2017.01.009

Yates, T. S., Fel, J., Choi, D., Trach, J. E., Behm, L., Ellis, C. T., & Turk-Browne, N. B. (2025). Hippocampal encoding of memories in human infants. Science, 387(6740), 1316–1320. 10.1126/science.adt7570

Yushkevich, P. A., Piven, J., Hazlett, H. C., Smith, R. G., Ho, S., Gee, J. C., & Gerig, G. (2006). User-guided 3D active contour segmentation of anatomical structures: Significantly improved efficiency and reliability. NeuroImage, 31(3), 1116–1128. 10.1016/j.neuroimage.2006.01.015

Yushkevich, P. A., Pluta, J. B., Wang, H., Xie, L., Ding, S.-L., Gertje, E. C., Mancuso, L., Kliot, D., Das, S. R., & Wolk, D. A. (2015). Automated volumetry and regional thickness analysis of hippocampal subfields and medial temporal cortical structures in mild cognitive impairment. Human Brain Mapping, 36(1), 258–287. 10.1002/hbm.22627

Zheng, A., Montez, D. F., Marek, S., Gilmore, A. W., Newbold, D. J., Laumann, T. O., Kay, B. P., Seider, N. A., Van, A. N., Hampton, J. M., Alexopoulos, D., Schlaggar, B. L., Sylvester, C. M., Greene, D. J., Shimony, J. S., Nelson, S. M., Wig, G. S., Gratton, C., McDermott, K. B., … Dosenbach, N. U. F. (2021). Parallel hippocampal-parietal circuits for self- and goal-oriented processing. Proceedings of the National Academy of Sciences, 118(34), e2101743118. 10.1073/pnas.2101743118

