## Supplemental_Materials for "Trajectories of hippocampal subregion development in the first years of life and their association with school-aged episodic memory outcomes"

Supplemental Analyses

*MST Recognition Score*

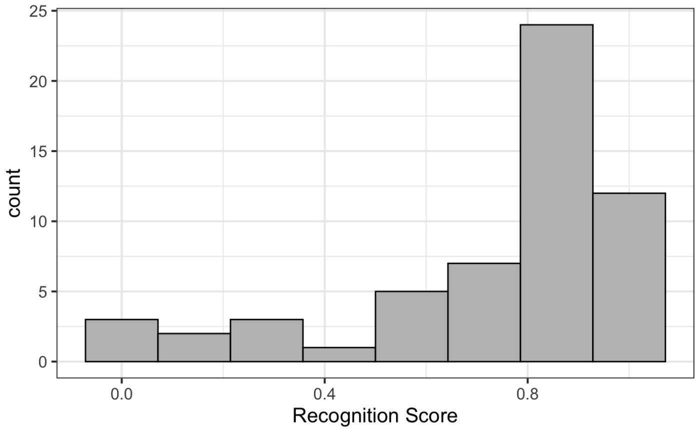


Supplemental Figure 1. Performance on the recognition score metric. 0.4 was used as a cut-off for sensitivity analyses.

*ASHS vs. nnUnet Hippocampal Subregion Segmentations*

Training data was also used as input for a new atlas in a preexisting hippocampal segmentation software, ASHS (Yushkevich et al., 2015). A new atlas was built based on the training set of manually segmented images following the procedures on the ASHS documentation site (https://sites.google.com/view/ashs-dox/local-ashs/building-an-atlas-for-t2-mri). We included a study-specific configuration file as the T2 images were isotropic, full-brain images rather than the MTL-specific T2 images expected by the standard ASHS software. Model performance was compared between ASHS and nnUnet qualitatively using visual review and quantitatively by calculating dice scores to evaluate segmentation overlap with the manual versions of the segmentations.

ASHS segmentations varied more widely than nnUnet segmentations. Four out of the 55 test set participants failed the segmentation pipeline altogether such that no segmentations were created. Of the 51 that completed, there were four participants with very low-quality segmentations (dice scores <0.6 even in the head or body of the hippocampus when compared to the gold standard manual segmentation), with two having dice scores of 0 for some regions, suggesting that the pipeline wasn't even able to identify the hippocampal subregion at all. Upon visual inspection, in general nnUnet segmentations looked more accurate than ASHS segmentations (ex. Supplemental Figure 2). Compared to the nnUnet segmentation, dice scores also showed a much stronger correlation with age (Supplemental Figure 3), suggesting that the ASHS pipeline is more biased by age. ASHS dice scores for all subregions showed a significant correlation with age (all $p<0.036$). The mean dice scores across subregions were also slightly lower than for nnUnet, with a mean between 0.82-0.85 across the head and body subregions, which are still relatively good, but below the nnUnet dice scores. The left and right tail subregions had a mean of 0.69 and 0.70 which is much lower than nnUnet and lower than what might be considered an acceptable average. Thus, nnUnet was selected as the final hippocampal subregion segmentation model.

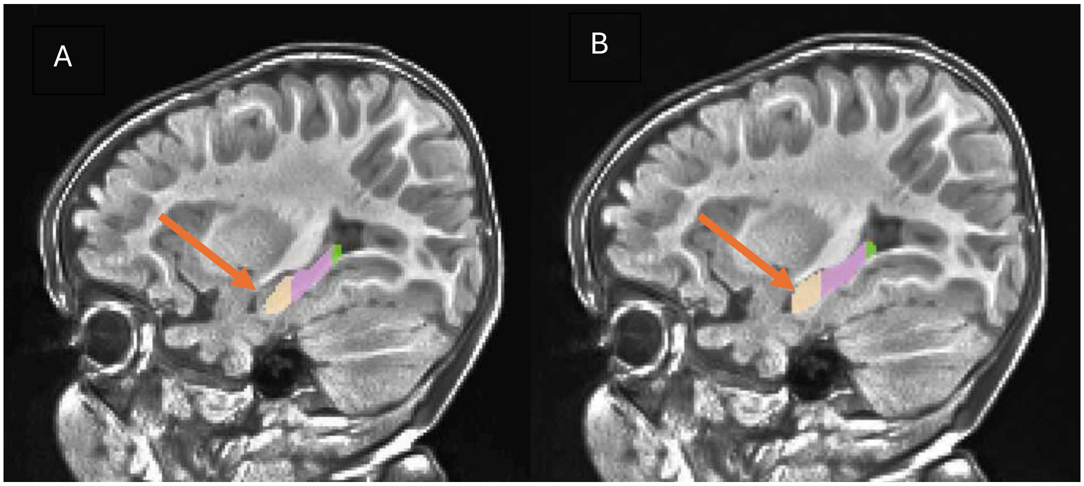


Supplemental Figure 2. Example hippocampal segmentation of a 12-month participant. A) ASHS segmentation missed a significant portion of the head of the hippocampus relative to the B) nnUnet segmentation

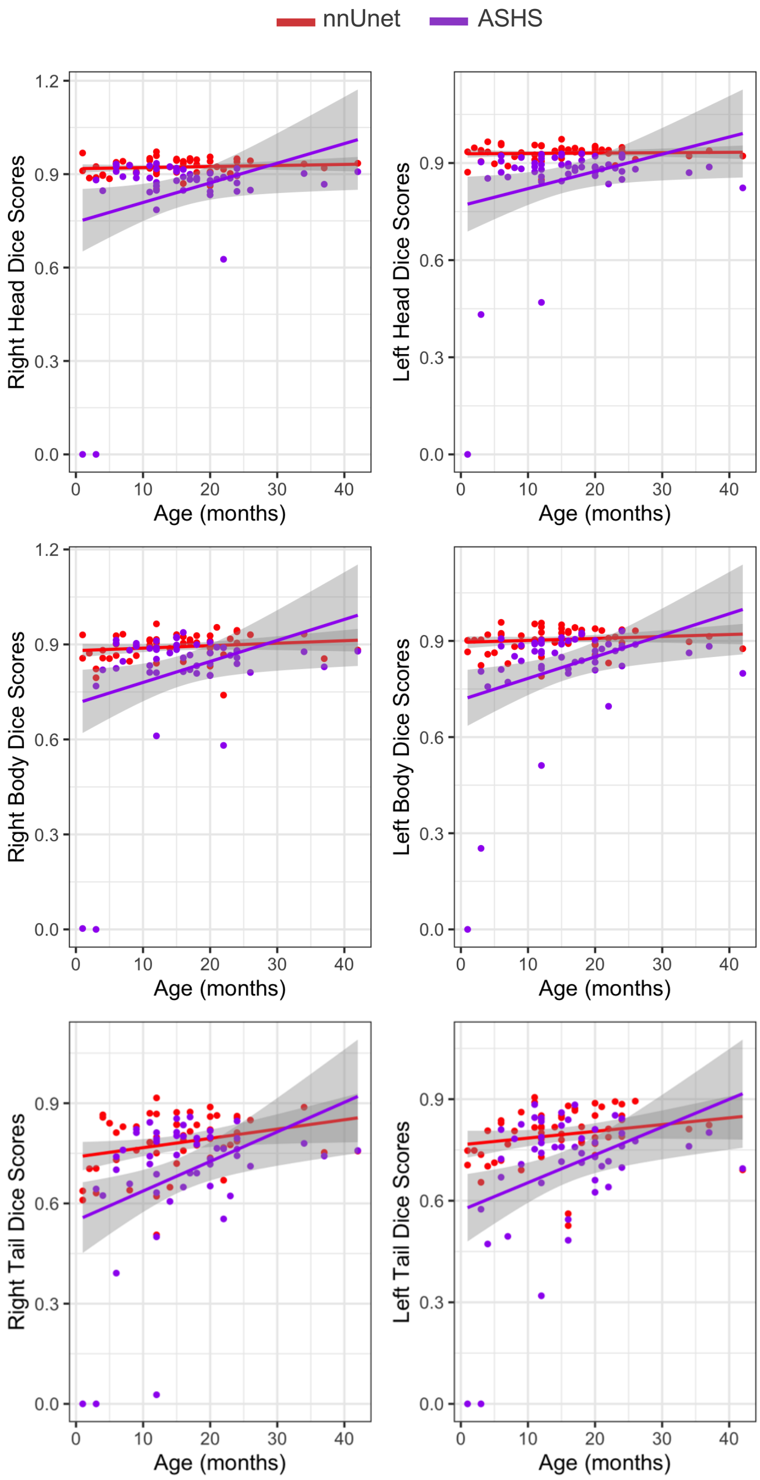


Supplemental Figure 3. Dice scores of the overlap between manual and automatic segmentations. Shown by age for the test set. Red=nnUnet; Purple=ASHS.

*Raw Data Demonstrates Hippocampal Head and Thalamus Hemisphere Differences*

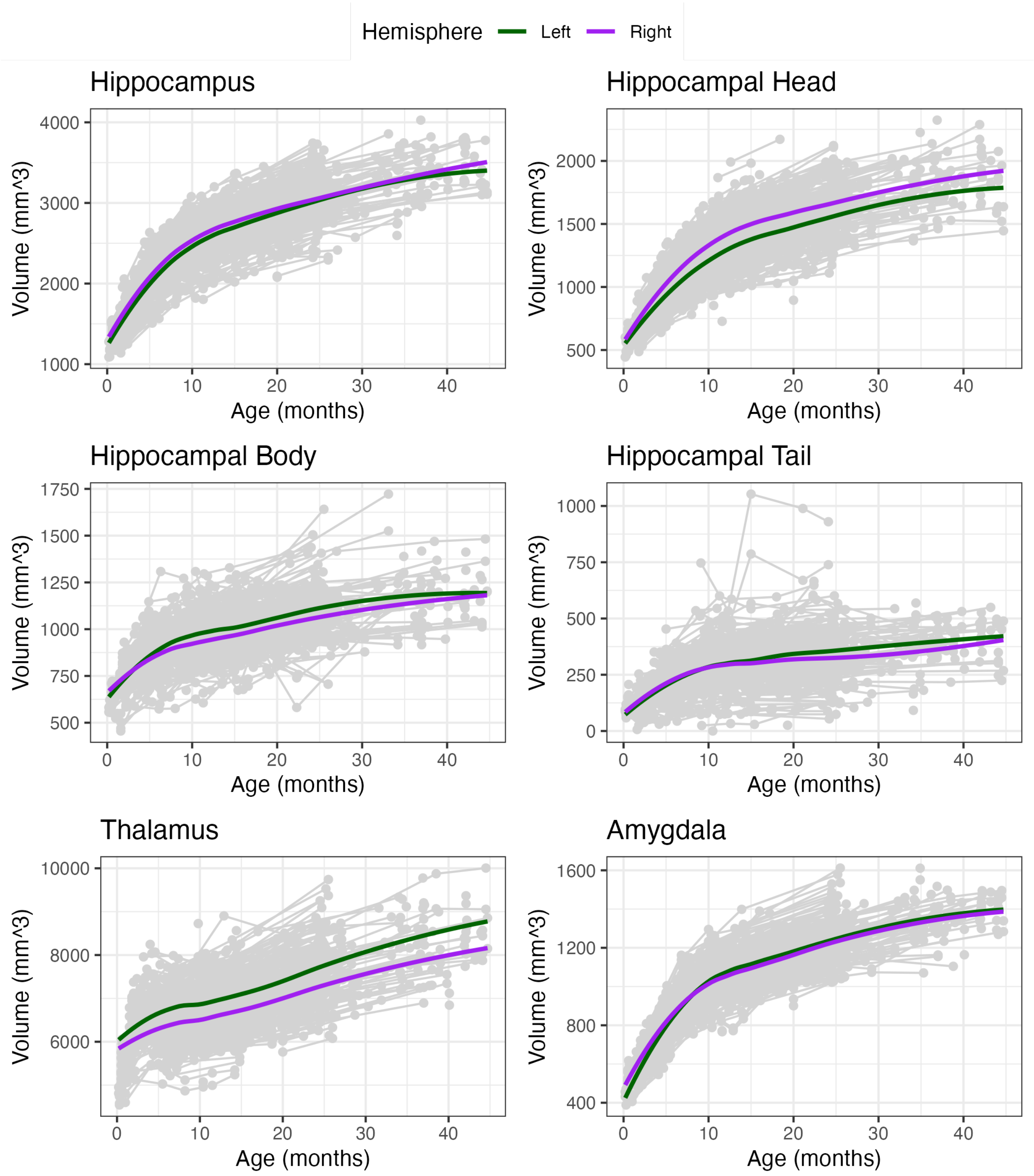


Supplemental Figure 4. Trajectories of subcortical regions associated with memory networks. The hippocampus was divided into subregions, and all regions are shown separated by left and right hemisphere. Smoothed loess curves are depicted on each plot. The head of the hippocampus and the thalamus both showed hemispheric differences in trajectory and so were separated for further analyses.

*Thalamus and Amygdala GAMM Trajectories*

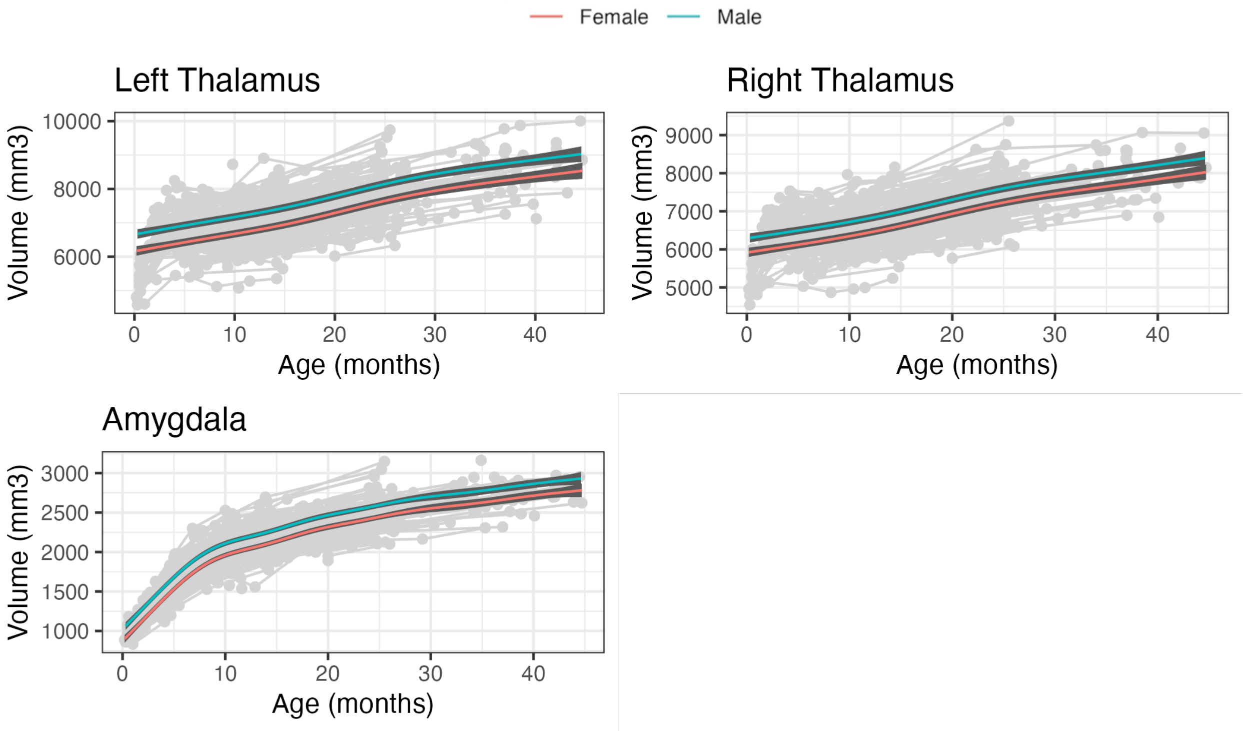


Supplemental Figure 5. Thalamus and amygdala regions were included in analyses as additional subcortical regions implicated in memory.

*Participant average residual*

The main statistical approach may represent a slight overestimation of the model, as mixed effects models including a random effect of participant did not converge, and so a sensitivity analysis was done using residuals averaged by participant. For participants with more than one visit, the difference in residuals between visits was visualized to determine the stability of the individual's relative trajectory. The residuals did seem to vary by age within a participant (Supplemental Figure 6B). Regardless, in the sensitivity analysis, the participant averaged residual was used as the predictor in the model, while still accounting for ages and the number of visits per participant. This model likely highly underestimates any effects due to averaging across meaningful information, but is provided as a sensitivity analysis for a more conservative estimate. No significant relationships were found using the participant averaged residuals across either the MST or source memory tasks.

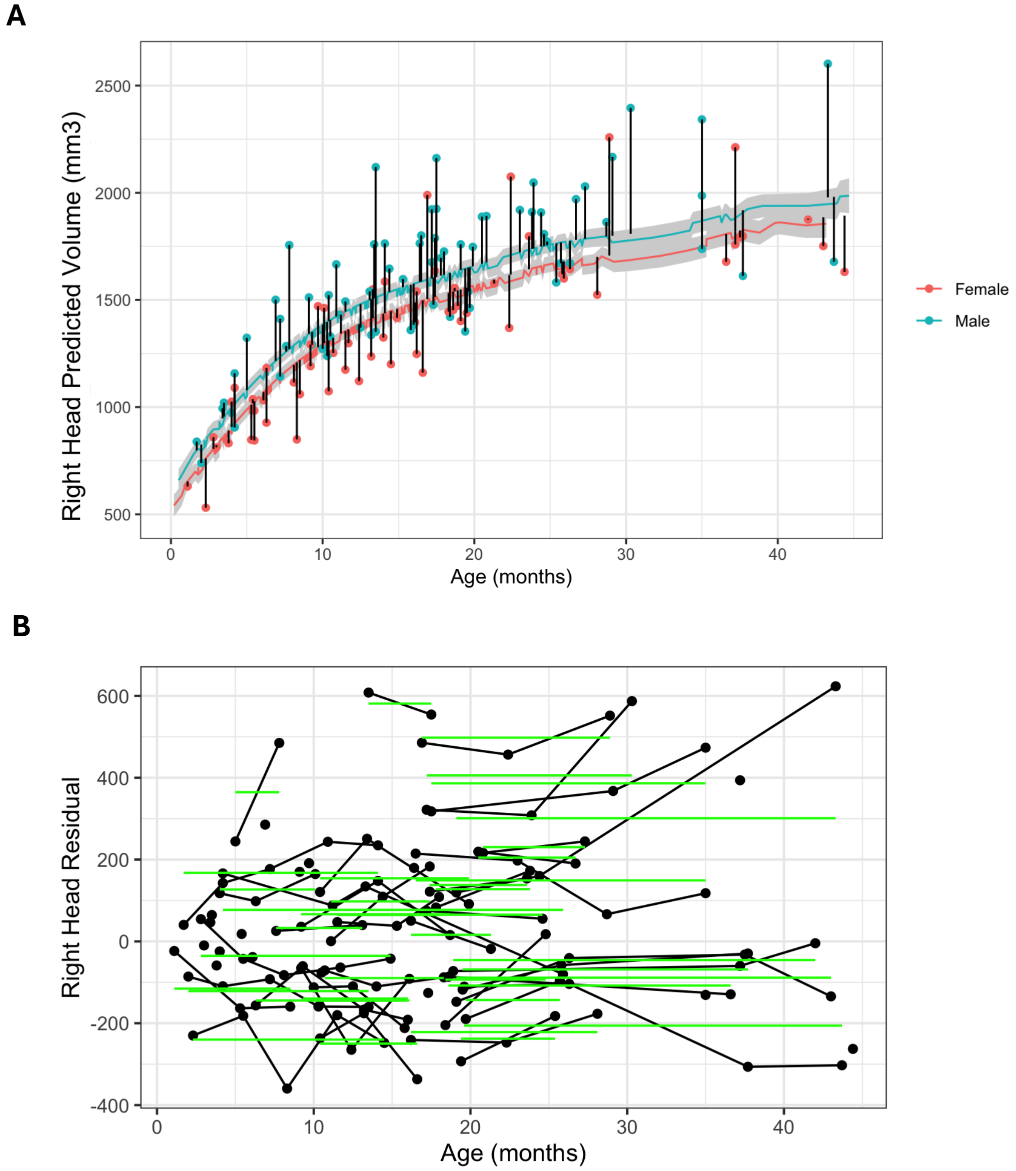


Supplemental Figure 6. GAMMs residual examples. A) An example of the residuals for the right head of the hippocampus for each of the visits for participants with episodic memory outcomes. B) An example of the residuals for the right head of the hippocampus plotted by age and participant. The horizontal lines represent each participant's average across visits, which suggests that an average across a participant might not be an ideal approach.

*nnUnet segmentation vs. manual behavioral differences*

Additionally, a sensitivity analysis was conducted using the hippocampal regions automatically calculated from nnUnet as opposed to manual segmentations, as both were available for these participants since they had been excluded from the training and testing sets during model training. For these participants, the mean difference between nnUnet vs. manual segmentation was 3% or less of each brain structure.

In these analyses, similar yet slightly different results were seen. GAMMs residuals of the right head of the hippocampus still showed a positive relationship with pattern discrimination performance on the MST, while the body of the hippocampus also showed this relationship with the nnUnet segmentations ($F(10, 120)=4.44$, $p_{Rhippohead}=0.003$, $p_{body}=0.041$). When using the GAMMs residuals that included an effect of total brain volume, the right head of the hippocampus, the body of the hippocampus, the right thalamus, and the amygdala all showed a significant relationship, with the right head and body of the hippocampus still positively predicting performance, and the thalamus (and now the amygdala) showing a negative relationship ($F(10, 120)=5.22$, $p_{Rhippohead}=0.021$, $p_{body}=0.008$, $p_{Rthalamus}=0.029$, $p_{amygdala}=0.044$). All of these relationships except the amygdala held after including the effect of FSIQ from the WPPSI-IV ($F(11, 117)=4.69$, $p_{Rhippohead}=0.010$, $p_{body}=0.012$, $p_{Rthalamus}=0.026$).

*MST alternative indices*

Three other indices were also calculated as described in Supplemental Table 1. The different indices represent different weightings of patterns of bias a participant might demonstrate. Sensitivity analyses explored relationships with brain development for each of these indices. The main relationships partially held across the other indices. No relationships were found with the “Target_FalseTarget” or “Lure_FalseLure” indices. In the “LureFoil_FalseTarget” index that was created specifically for these participants, the body of the hippocampus and left and right thalamus all showed relationships ($F(10, 120)=4.18$, $p_{body}=0.046$, $p_{Lthalamus}=0.016$, $p_{Rthalamus}=0.003$), but the head of the hippocampus did not remain significant.

Supplemental Table 1. Indices of Pattern Discrimination

| Citation | Index Abbreviation | Formula^ | Description |
|---|---|---|---|
| (Ngo et al., 2018) | Lure_FalseTarget | similar\|similar - same\|similar | Main outcome of interest |
| (Canada et al., 2019) | Target_FalseTarget | same\|same – same\|similar | Another index used in MST literature in this age that focuses on differences in discrimination performance and those correctly assigned same instead of those correctly assigned similar |
| (Stark et al., 2019) | Lure_FalseLure | similar\|similar – similar\|new | Classic index used in adult MST literature |
| n/a | LureFoil_FalseTarget | (new or similar)\|(new or similar) – same\|(new or similar) | New index that combines new and similar images in order to capture a subset of participants who didn't endorse any images as similar |

^Formula represents response|actual, where similar|similar would be the proportion of objects where children accurately assigned "similar" to similar objects. Same|similar would represent the proportion of images where children incorrectly assigned "exactly the same" to similar objects.
